# Rescuing perishable neuroanatomical information from a threatened biodiversity hotspot: Remote field methods for brain tissue preservation validated by cytoarchitectonic analysis, immunohistochemistry, and X-ray microcomputed tomography

**DOI:** 10.1101/052415

**Authors:** Daniel F. Hughes, Ellen M. Walker, Paul M. Gignac, Anais Martinez, Kenichiro Negishi, Carl S. Lieb, Eli Greenbaum, Arshad M. Khan

## Abstract

Biodiversity hotspots, which harbor more endemic species than elsewhere on Earth, are increasingly threatened. There is a need to accelerate collection efforts in these regions before threatened or endangered species become extinct. The diverse geographical, ecological, genetic, morphological, and behavioral data generated from the on-site collection of an individual specimen are useful for many scientific purposes. However, traditional methods for specimen preparation in the field do not permit researchers to retrieve neuroanatomical data, disregarding potentially useful data for increasing our understanding of brain diversity. These data have helped clarify brain evolution, deciphered relationships between structure and function, and revealed constraints and selective pressures that provide context about the evolution of complex behavior. Here, we report our field-testing of two commonly used laboratory-based techniques for brain preservation while on a collecting expedition in the Congo Basin and Albertine Rift, two poorly known regions associated with the Eastern Afromontane biodiversity hotspot.

First, we found that transcardial perfusion fixation and long-term brain storage, conducted in remote field conditions with no access to cold storage laboratory equipment, had no observable impact on cytoarchitectural features of lizard brain tissue when compared to lizard brain tissue processed under laboratory conditions. Second, field-perfused brain tissue subjected to prolonged post-fixation remained readily compatible with subsequent immunohistochemical detection of neural antigens, with immunostaining that was comparable to that of laboratory-perfused brain tissue. Third, immersion-fixation of lizard brains, prepared under identical environmental conditions, was readily compatible with subsequent iodine-enhanced X-ray microcomputed tomography, which facilitated the non-destructive imaging of the intact brain within its skull.

In summary, we have validated multiple approaches to preserving intact lizard brains in remote field conditions with limited access to supplies and a high degree of environmental exposure. This protocol should serve as a malleable framework for researchers attempting to rescue perishable and irreplaceable morphological and molecular data from regions of disappearing biodiversity. Our approach can be harnessed to extend the numbers of species being actively studied by the neuroscience community, by reducing some of the difficulty associated with acquiring brains of animal species that are not readily available in captivity.

## 1. Introduction

By one estimate [1], 86% of the world’s extant eukaryotic species still await identification and description. It is believed that our current classification and taxonomic efforts are too slow to overcome biodiversity loss [1]. As a result, many species may go extinct before their existence is even known to us. Terrestrial biodiversity is concentrated in at least 35 biodiversity hotspots. Although they account for only 2.3% of the Earth’s land surface, these areas harbor over 50% of the world’s endemic plant species and an estimated 43% of endemic terrestrial vertebrate species [2]. Intensive efforts are now underway to fully characterize and document the biota within these hotspots, which are anticipated to yield the highest amount of data in the shortest amount of time [3]. Thus, even if rapid global biodiversity loss cannot be fully prevented, efforts can be made at these hotspots to mitigate data losses with targeted efforts at data rescue. Such efforts can help inform rational strategies for conservation efforts that have been demonstrated to slow the rate of global biodiversity decline [4] and help increase our understanding of how traits vary across species.

An important part of such data rescue involves documenting biodiversity through the careful and responsible on-site collection of individual members of poorly known species [5, 6]. On-site collection allows for a variety of information to be gathered for such species, including geographical, ecological, genetic, biochemical, morphological, and behavioral datasets [e.g., 7–11]. Having diverse datasets for a species, in turn, affords investigators flexibility in how the data can later be used for a host of analytical approaches across molecular to macro-evolutionary scales [12–19], even if current paradigms of analysis favor some datasets over others.

A potentially useful—but often overlooked—source of variation is the brain. Mapping of neuroanatomical characters onto molecular-based phylogenies has revealed new information about differences in brain region size and encephalization among species [14–16], the evolution of species-specific communication [17], and the evolutionary origins of the neurological configuration of the brain for certain taxa [18]. Moreover, comparing neuroanatomical characters in wild-caught animals with those in their domesticated counterparts has provided insights about the genetic routes through which domestication becomes manifest in different species [19]. Unfortunately, field methods used to preserve collected specimens traditionally have been incompatible with the preservation of neuroanatomy for several reasons. First, biodiversity hotspots are often located in remote regions of developing countries where infrastructure is inadequate to support laboratory-based neuroanatomical and neuromolecular research [20, 21]. Second, in remote field locations there is limited access to resources that ensure optimal preservation of brain tissue, such as appropriate fixatives or stable cold storage conditions free from environmental exposure. Finally, local regulations often prohibit the export of live animals to outside countries where adequate laboratory-based infrastructure may exist, further discouraging researchers without access to regional laboratories from preserving brain tissues optimally.

In addition to these challenges, the collection of rare and previously undocumented species affords additional considerations related to brain tissue processing. In particular, dissecting and sectioning preserved brains destroys important gross anatomical information that is potentially useful for advancing knowledge of the brains of newly discovered or previously undocumented species. Such information includes craniometric relationships between the skull and underlying brain tissue structures that could potentially inform future stereotaxic procedures, as well as three-dimensional relationships within the brain and between cranially derived sensory and motor organs and the neural networks to which they are connected. This information in turn can help enable classification of cell types, neural configurations, and structure-function relationships for specific brain circuits across a far wider diversity of vertebrate taxa than is currently understood.

In this study, portions of which have been presented in preliminary form [22], we have developed a validated field protocol for brain tissue preservation that overcomes these challenges. Specifically, two of us (DFH and EG) undertook a 58-day collecting expedition to the Congo Basin and Albertine Rift of Central Africa, both poorly known regions that form portions of the Eastern Afromontane biodiversity hotspot [23, 24], and performed on-site euthanasia of lizards and fixation of their intact brains under entirely remote conditions, with limited access to supplies, and during a high degree of environmental exposure. We field-tested two tissue fixation methods commonly used in the laboratory: immersion and transcardial perfusion with buffered formalin. We evaluated the efficacy of these methods in the laboratory by examining the field-fixed samples collected in Central Africa at the cytoarchitectural, chemoarchitectural, and gross-neuroanatomical levels by using semi-quantitative Nissl-based structural analysis, immunohistochemistry, and diffusible iodine-based contrast-enhanced computed tomography (diceCT) [25, 26], respectively. Our field protocols not only generated high-quality tissue preservation at the cellular and regional tissue levels, but they are also compatible with non-destructive imaging, at the gross neuroanatomical level, of the intact skull and underlying soft brain tissue. These fixation methods are simple to implement in the field, require few resources that would otherwise be diiffcult to obtain in remote locations, and are extensible to collection efforts for a variety of poorly known or undiscovered vertebrates found in the world’s most fragile ecosystems.

## 2. Materials and Methods

### 2.1 Approvals and permissions

Permission to collect lizards in Uganda was obtained from the Uganda Wildlife Authority (UWA), the National Biodiversity Data Bank at Makerere University, Institut Superieur d’Ecologie pour la Conservation de la Nature (ISEC), and Uganda’s CITES License (2888). Permission to collect in Democratic Republic of Congo (DRC) was granted by the Centre de Recherche en Sciences Naturelles (CRSN – LW1/27/BB/KB/BBY/60/2014) and the Institut Congolais pour la Conservation de la Nature (ICCN – 1007/ICCN/DG/ADG/DT/04). The University of Texas at El Paso’s (UTEP) Institutional Animal Care and Use Committee (IACUC – A-200902-1) approved field and laboratory methods.

### 2.2 Expedition details and experimental subjects collected for this study

The expedition took place May–July 2014. **Table 1** lists the animals collected for this study, including the locations where they were collected. Location 1 is in Bwindi Impenetrable National Park, Uganda; Location 2 is in Rwenzori Mountains National Park, Uganda; and Location 3 is in the small village of Boda in northeastern DRC. These collection sites are located in the Albertine Rift (Locations 1, 2) and Congo Basin (Location 3), two regions that form portions of the Eastern Afromontane biodiversity hotspot [23, 24]. In addition to animals collected in the field, control animals listed in **Table 2** and processed at UTEP under laboratory conditions were purchased from Underground Reptiles (Deerfield Beach, FL).

**Table 1.** Details for the animals collected in the field

| Species | ID | SVL | TL | BW | Sex | Coordinates | Elevation | Fixation Type, Time* | Duration w/o Cold Storage |
| --- | --- | --- | --- | --- | --- | --- | --- | --- | --- |
| **Location 1: Uganda: Western Region, Kabale-Kanungu Districts, Bwindi Impenetrable National Park** |  |  |  |  |  |  |  |  |  |
| <u>*Trioceros johnstoni*</u><br>[see Figs. 2 & 6A] | UTEP 21385 | 109 | 121 | 31.9 | M | S01.04836,<br>E29.77684 | 2284 m | P, 58 min | 54 d |
| *Rhampholeon boulengeri* | UTEP 21386 | 48 | 13 | 2.5 | Jv, M | S00.97828,<br>E29.69354 | 1563 m | P, 42 min | 54 d |
| **Location 2: Uganda: Western Region, Kasese District, Rwenzori Mountains National Park** |  |  |  |  |  |  |  |  |  |
| *Trioceros ellioti* | UTEP 21387 | 59 | 53 | 3.9 | M | N00.34972,<br>E30.02973 | 1655 m | P, 51 min | 51 d |
| <u>*Trioceros johnstoni*</u><br>[see Fig. 7B,C] | UTEP 21388 | 100 | 107 | 31.2 | F | N00.36033,<br>E30.00975 | 1909 m | I, 18 min | NA |
| <u>*Rhampholeon boulengeri*</u><br>[see Fig. 7A] | UTEP 21389 | 47 | 12 | 3.6 | F | N00.36029,<br>E30.00922 | 1942 m | I, 25 min | NA |
| *Rhampholeon boulengeri*<br>[see Fig. 6C] | UTEP 21390 | 46 | 14 | 3.1 | M | N00.36029,<br>E30.00922 | 1942 m | P, 33 min | 51 d |
| **Location 3: DR Congo: Orientale Province, Bas-Uele District, Boda village** |  |  |  |  |  |  |  |  |  |
| <u>*Agama cf. finchi*</u><br>[see Fig. 4] | UTEP 21391 | 108 | 171 | 40.9 | M | N03.52319,<br>E26.39019 | 653 m | P, 46 min | 21 d |

Details regarding the animals which underwent euthanasia and fixation under field conditions for this study. Abbreviations used: BW, body weight (g); F, female; I, immersion fixation; ID, identification number (UTEP Biodiversity Collections); Jv, juvenile; M, male; NA, not applicable (these specimens did not go to cold storage); P, perfusion fixation; SVL, snout–vent length (mm); TL, tail length (mm). Note that coordinates are expressed in decimal degrees. Specimens indicated by underlining are those for which tissue photographs have been furnished in this study (brackets note the specific figures). \* The interval between sedation of the subject to storage of the fixed brain.

**Table 2.** Details for the animals used in the laboratory

| Species | ID | SVL | TL | BW | Sex | Fixation Type | Time* | Duration w/o Cold Storage |
| --- | --- | --- | --- | --- | --- | --- | --- | --- |
| <u>*Trioceros jacksonii*</u><br>[see Fig. 3] | UTEP 21382 | 111 | 92 | 40.9 | F | P | 34 min | 0 d |
| *Trioceros jacksonii* | UTEP 21383 | 119 | 114 | 43.5 | M | P | 50 min | 0 d |
| <u>*Rieppeleon kerstenii*</u><br>[see Fig. 6B,D] | UTEP 21384 | 49 | 10 | 3.2 | F | P | 29 min | 0 d |

### 2.3 Formaldehyde sources

#### 2.3.1 Sources used for field studies

The buffered formalin solution used in this study was derived from either of two sources: (1) stock formalin (37% v/v formaldehyde) sold commercially in bottled liquid form in Kampala, Uganda and (2) paraformaldehyde powder (100% formaldehyde) obtained from the University of Kisangani, DRC. These sources of formaldehyde were used to freshly prepare 1 L batches of ca. 4% and 10% buffered formalin (100 ml v/v of source (1) or 100 g w/v of source (2) added to 900 ml of water) in the field using 4 g of sodium phosphate monohydrate (NaH_2_PO_4_H_2_O) and 6.5 g of dibasic sodium phosphate anhydrate (Na_2_HPO_4_) per liter of formalin. The solution prepared from powdered formaldehyde differed from the liquid commercial-grade formalin in a few important ways: (1) it was mixed without heat-mediated depolymerization, due to a lack of electricity and laboratory facilities; (2) it was not filtered; (3) it contained a final concentration of 10% formaldehyde; and (4) it lacked methanol, a common stabilizer which can affect certain immunological reactions. In order to facilitate semi-quantitative comparisons with laboratory-fixed chameleons (fixed using formalin with a final concentration of 4% formaldehyde), the field-caught chameleons listed in Table 1 (Locations 1 and 2) were all fixed using the diluted liquid stock solution (ca. 4% formaldehyde, final concentration). In contrast, the agamid (Location 3) was the only specimen perfused using the solution prepared from powdered fixative (10% formaldehyde, final concentration). Although the pH values for these solutions were not measured in the field, they were likely near neutral pH, based on the buffering ranges of the salts we used.

#### 2.3.2 Source used for laboratory studies

The fixative used was prepared from freshly depolymerized and cleared granular *p*-formaldehyde (Electron Microscopy Sciences Inc., Hatfield, PA; Catalog #19210) as a 4% w/v solution in sodium borate buffer (pH 9.5 at 4°C). First validated by Berod and colleagues [27], this high pH solution is used routinely in our laboratory for locating neural antigens with immunohistochemistry [28-31].

## 2.4 Methods pertaining to tissue fixed by transcardial perfusion

### 2.4.1 Transcardial perfusions

#### 2.4.1a: Transcardial perfusions under field conditions

**Figure 1** shows details of our field procedures, and **Table 3** lists the supplies used to perform them. Lizards were deeply sedated by placing them in closed plastic containers containing two cotton balls saturated with liquid isoflurane. When the animals were sedated enough to remain immobile, they were briefly removed from the container to record body weight and snout-vent length before being returned to the container to complete the sedation. Other more detailed morphometric measurements necessary for biodiversity studies, especially of the head, were also recorded at this time (e.g., head length and width, snout length, etc.). Once fully anesthetized (i.e., no Labyrinthine righting reflex), animals were affixed to a silicone mat by a single pin pierced through each appendage (**Fig. 1B**).

**Figure 1.**
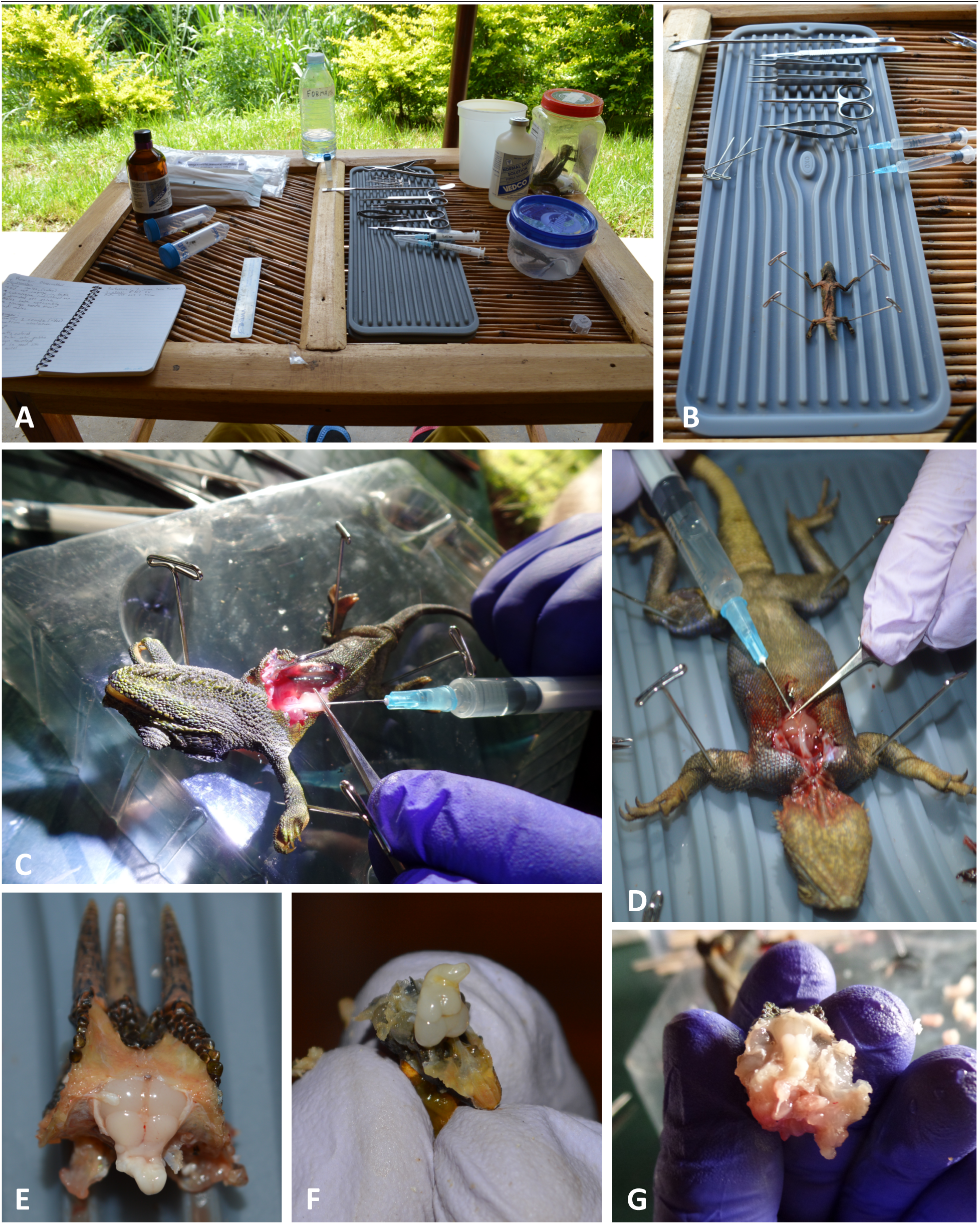
Images of field-based perfusion technique. The field laboratory setup (A); pinned lizard on silicone mat prior to opening of the thoracic cavity (B); injections of solution through opening in apex of heart (C, D); partially dissected and exposed formaldehyde-fixed brains (E, F, G).

**Table 3.** Materials for field perfusion procedures

| Item | Quantity | Supplier | Catalog # |
| --- | --- | --- | --- |
| **1. Containers** |  |  |  |
| Falcon™ conical centrifuge tube (polypropylene, 50 ml) | 5 | Fisher | 352070 |
| field box with handle (11.6" x 5.1" x 7.1") | 1 | Plano Molding | 131200 |
| Fisherbrand™ bottle (polyethylene, 125 ml) | 10 | Fisher | 02911952 |
| **2. Reagents and solutions** |  |  |  |
| 4% and 10% formalin, sodium phosphate buffered | varied | *See Methods* | NA |
| Isosol™ (isoflurane, USP) (250 ml) | 2 | Vedco | NDC 50989-150-15 |
| normal saline solution, sterile (250 ml) | 2 | Vedco | NDC 50989-641-15 |
| sucrose (5 kg) | 30 x 3 g | Sigma-Aldrich | S8501 |
| **3. Perfusion and dissecting instruments** |  |  |  |
| hypodermic needle (18 ga) | 10 | Nipro | AH+1825 |
| syringe (3 cc) | 3 | Nipro | JD+03L |
| Dumont #5SF forceps (inox steel, super fine, straight tip) | 1 | Fine Science Tools | 11252-00 |
| Friedman-Pearson rongeurs (1 mm cup size, straight tip) | 1 | Fine Science Tools | 16020-14 |
| interchangeable blades (angled, 10 mm cutting edge) | 10 | Fine Science Tools | 10035-15 |
| Moria fine scissors (inox steel, extra sharp, straight tip) | 1 | Fine Science Tools | 14370-22 |
| insect pin (size 3, 0.5 mm diameter, 4 cm length) | 10 | Fine Science Tools | 26001-50 |
| scalpel handle #3 (stainless steel, 12 cm length) | 1 | Fine Science Tools | 10003-12 |
| spatula & probe (stainless steel, 14 cm length) | 1 | Fine Science Tools | 10090-13 |
| student surgical scissors (stainless steel, 14.5 cm length) | 1 | Fine Science Tools | 91402-14 |
| Vannas spring scissors (straight tip, 2 mm cutting edge) | 1 | Fine Science Tools | 15000-03 |
| **4. Miscellaneous supplies** |  |  |  |
| cotton ball (500/pack) | 1 | U.S. Cotton |  |
| digital balance, battery-operated | 1 | Ohaus |  |
| Parafilm "M" (2" x 250') | 10 (strips) | Bemis | PM992 |
| silicone mat | 2 | OXO | 372100V2 |
| plastic ruler | 1 |  |  |

List of materials used for the field perfusion procedures.

For perfusion, the lizard’s snout was placed inside a 50-ml conical tube that contained an isoflurane-soaked cotton ball at its base. To open the mediastinum (thoracic cavity), scissors, aided with finer incisions from a scalpel, were used to cut anterior-posterior from mid-neck to lower-abdomen. The thoracic cavity was opened without collapsing the pectoral girdle on the brachiocephalic trunks and carotid arteries. The thoracic wall was removed—the sternum was cut free from the ribs, connective tissue was excised, and portions of the ribs and lungs were removed to further expose the heart, right atrium, and carotid arteries (**Fig. 1C and D**). The common carotid artery was gently seized with forceps to elevate the heart from the pericardial cavity and better observe the flow of injected solutions toward the head (**Fig. 1C and D**). Lizards were exsanguinated from an incision to the right atrium with fine scissors. Two 3-ml syringes, equipped with 18-gauge needles, were used for successive injections into the apex of the heart (**Fig. 1C and D**). The needle tip was inserted carefully into the apex and extended through the ventricle to settle visibly just beyond the base of the common carotid. Saline was injected first, followed by buffered formalin solution. In both cases, due to lack of ice or cold storage, the solutions injected were not cold. The small amount of liquid formalin waste (ca. 2 ml) collected after perfusion was diluted with water to a nonhazardous concentration of < 0.1% and disposed of down a drain.

#### 2.4.1b: Transcardial perfusions under laboratory conditions

Control-group animals (**Table 2**) perfused transcardially under laboratory conditions at UTEP underwent identical procedures to those described above for field perfusions with a few notable differences. First, the formulation and source of formaldehyde used were different than the sources used in the field (see *Section 2.3.2*). Second, when saline and fixative were successively injected into the animal, both solutions were ice cold. Finally, perfusions were performed in a chemical fume hood.

### 2.4.2 Brain dissection

#### 2.4.2a: Dissections in the field

The head of the euthanized and perfused animal was removed above the shoulders with large surgical scissors. If still attached, cervical vertebrae were removed with rongeurs. To uncover the occipital and parietal skull bones, the postcranial musculature and surrounding connective tissue were gently removed with fine scissors or scraped away by scalpel. To facilitate manipulation of the cranium, the lower mandible was separated from the head with rongeurs. Dorsal portions of the parietal and temporal skull bones were removed, and the entire occipital skull bone was excised. Saline irrigation helped to maintain moisture levels in the brain tissue once it was exposed to the environment. All connective tissues between the skull and brain (i.e., meninges) were gently teased apart, and the roots of the cranial nerves severed, thereby releasing the brain from its remaining attachments to the skull. The unattached brain was removed from the cranial cavity and placed immediately into storage solution (see *Section 2.4.3a*).

#### 2.4.2b: Dissections in the laboratory

One major difference between field-based dissections and those performed in the laboratory involved temperature control. Specifically, following transcardial perfusions in the laboratory, the heads were removed and placed immediately on ice. Fixed and chilled brains were excised a few hours later as described previously. After dissection, these brains were stored immediately in storage solution (*Section 2.4.3b*) at 4°C.

### 2.4.3 Brain storage

#### 2.4.3a: Storage conditions in the field

Following dissection, brains were stored in individually labeled 100 ml plastic vials filled with a buffered formalin solution containing 12% w/v sucrose (“storage solution”) [28-31]. The solution was topped off to minimize evaporative loss and to ensure that the brain would be wholly submerged. Infiltration of sucrose was confirmed when each brain lost buoyancy and sank to the bottom of the vial. Liquid levels in the vials were checked daily and replenished if low. Care was taken to avoid exposing the vials to excessive heat. Upon arrival at the UTEP Systems Neuroscience Laboratory, these brains were placed initially in a cold room (4°C), and after a brief period (ca. 48 hours), the perfusion-fixed brains were frozen as described *(Section 2.4.4)*.

#### 2.4.3b: Storage conditions in the laboratory

Each brain remained in storage solution (the same fixative solution noted in *Section 2.3.2*, with 12% w/v sucrose [28-31]) at 4°C, until sinking to the bottom of its vial.

### 2.4.4 Freezing of brains and histology

The following procedures were conducted at the UTEP Systems Neuroscience Laboratory. Brains collected in the field or in the laboratory were removed from their respective storage solutions, blotted dry, and then flash frozen in a plastic container filled with hexane supercooled over a bed of powdered dry ice. The frozen brains were then stored at −80°C until further processing. To prepare them for sectioning, all brain samples were placed into small plastic molds, embedded in Tissue-Tek OCT embedding medium (10.24% polyvinyl alcohol, 4.26% polyethylene glycol, and 85.5% non-reactive ingredients; Sakura Finetek USA, Inc., Torrance, CA), and returned to the −80°C storage until the embedding medium hardened. The OCT medium helped to maintain tissue stability, especially for the smallest lizard brain samples, throughout the sectioning process. Each OCT-embedded brain block was cut into 20-30 μm-thick sections in the transverse (coronal) plane using a Reichert sliding microtome (Reichert Austria Nr. 15 156) fitted with a modified brass freezing stage (Brain Research Laboratories, Newton, MA; Cat #3488-RJ) chilled with powdered dry ice. Four serial series of brain sections were collected in 24-well plates filled with anti-freeze cryoprotectant solution (50% 0.1 M sodium phosphate buffer, 30% ethylene glycol, and 20% glycerol; [32]). Sections were maintained in cryoprotectant at –20°C until further processing.

### 2.4.5 Nissl staining

Freely floating sections were rinsed twice in an isotonic Tris-buffered saline (TBS; pH 7.6 at room temperature) to wash out cryoprotectant. Sections were mounted on gelatin-coated slides using a fine-tipped paintbrush. Mounted sections were dried overnight (24 h) at room temperature (ca. 20°C) in a vacuum chamber. They were then dehydrated in ascending ethanol concentrations (50–100%; 3 min each), defatted in xylene, stained in 0.5% w/v thionine solution (thionin acetate, Catalog #T7029; Sigma-Aldrich Corporation, St. Louis, MO) [33, 34], and differentiated in 0.4% anhydrous glacial acetic acid. Slides were coverslipped with DPX mounting medium (Catalog # 06522; Sigma-Aldrich) and stored flat within covered slide trays.

### 2.4.6 Photomicrography and post-acquisition image processing of Nissl-stained tissues

Nissl-stained tissues were examined under bright field illumination using a Zeiss M2 AxioImager microscope equipped with an X-Y-Z motorized stage (Carl Zeiss Corporation, Thornwood, NY). Wide field mosaic images of stained histological sections were obtained using a cooled EXi Blue camera (QImaging, Inc., Surrey, British Columbia, Canada) driven by Volocity Software (Version 6.1.1; Perkin-Elmer, Inc., Waltham, MA) installed on the Apple Mac Pro computer. Images were exported from Volocity as lossless TIFF formatted files and imported into Adobe Photoshop (Version CS6; Adobe Systems, Inc., San Jose, CA). In Photoshop, images were cropped via the lasso tool, converted and resampled to 300 dpi gray scale, and brightness-and contrast-adjusted via the curves tool. For hollow spaces within the tissue (e.g., third ventricle), any background illumination of the slide remaining after white balancing was not cropped. Care was taken to make all adjustments judiciously across photomicrographs of field-and lab-processed tissue sections and also to track changes in scale during any size conversion.

### 2.4.7 Semi-quantitative histological evaluation

To evaluate the relative efficacies of the lab-and field-based perfusion methods for tissues fixed with 4% formaldehyde (final concentration), the condition of the sections obtained by each procedure was evaluated using a semi-quantitative approach. Three members of the UTEP Systems Neuroscience Laboratory (EMW, AM, and KN), who were blind to the treatment conditions of the tissue, independently rated tissue sections from both treatments using a three-point quality scale: poor (1), good (2), and excellent (3). This rating was performed by viewing the tissue sections under bright field illumination. To understand the effect of perfusion treatment (laboratory vs. field) on various aspects of both tissue and stain quality, independent raters were instructed to apply this scale across six criteria: (A) presence of blood in the tissue; (B) evenness of stain; (C) integrity of tissue at the center of the section; (D) integrity of tissue at the edges of the section; (E) clarity of lamination patterns; and (F) visibility and clarity of nuclei and cell clusters. A total of 204 stained tissue sections were evaluated, representing 166 sections prepared under laboratory conditions and 38 sections prepared under field conditions. The sample size discrepancy between treatments was not believed to influence the overall results because the model-based statistical approach we used (see below) is a function of the size of the largest cluster rather than of the number of clusters (i.e., population-averaged estimates).

### 2.4.8 Statsitical analyses

A heat map of the scores obtained by our independent observers was generated using R function heat map [35]. Because criteria C-F (see *Section 2.4.7*) are not independent from criteria A and B (i.e., the criteria are interdependent) and because both fixed effects (field vs. lab conditions) and random effects (order of tissue sampling onto slides, variable sample thickness, and subject selected for perfusion fixation) were present, we employed a variation of a general linear mixed model to analyze the results. Specifically, to best compare the scores between the two fixation methods, a generalized estimating equation (GEE) approach [36] was used to account for the dependence structure among clustered ordinal scores, as implemented in *geepack* package [37] in R. We tested each variable separately. The analyses were conducted based on the raw data and a reduced dataset using the mode score for each slide from each observer. This data reduction did not influence the results.

### 2.4.9 Immunohistochemistry

To evaluate the effect of the perfusion method on the immunoreactivity of neural antigens in the tissue samples, we performed a series of indirect immunohistochemistry experiments using distinct primary antibodies (**Table 4**) across both field-and laboratory-processed tissue samples preserved using solutions containing 4% formaldehyde. In the first set of experiments, we incubated tissue sections from field-(*Trioceros johnstoni*) and laboratory-perfused *(Rieppeleon kerstenii)* chameleons with a rabbit polyclonal antibody targeting the catecholamine-synthesizing enzyme tyrosine hydroxylase (TH). Importantly, the specificity of this TH antibody has been validated in Western blots as recognizing a single, 62 kDa protein band from both amphibian and reptile brain homogenates and shown to be immunogenically identical to that of mammalian TH [38, 39]. In a second set of experiments, field-(*Rhampholeon boulengeri*) and laboratory-perfused *(Rieppeleon kerstenii)* chameleon tissue sections were incubated with an antibody targeting neuropeptide Y (NPY). In some initial experiments, we also co-incubated the TH antibody with an antibody targeting dopamine beta-hydroxylase (DβH), and the NPY antibody with an antibody targeting calbindin (**Table 4**). However, since these antibodies produced little to no specific staining, they were not used in subsequent immunohistochemical runs.

**Table 4.**
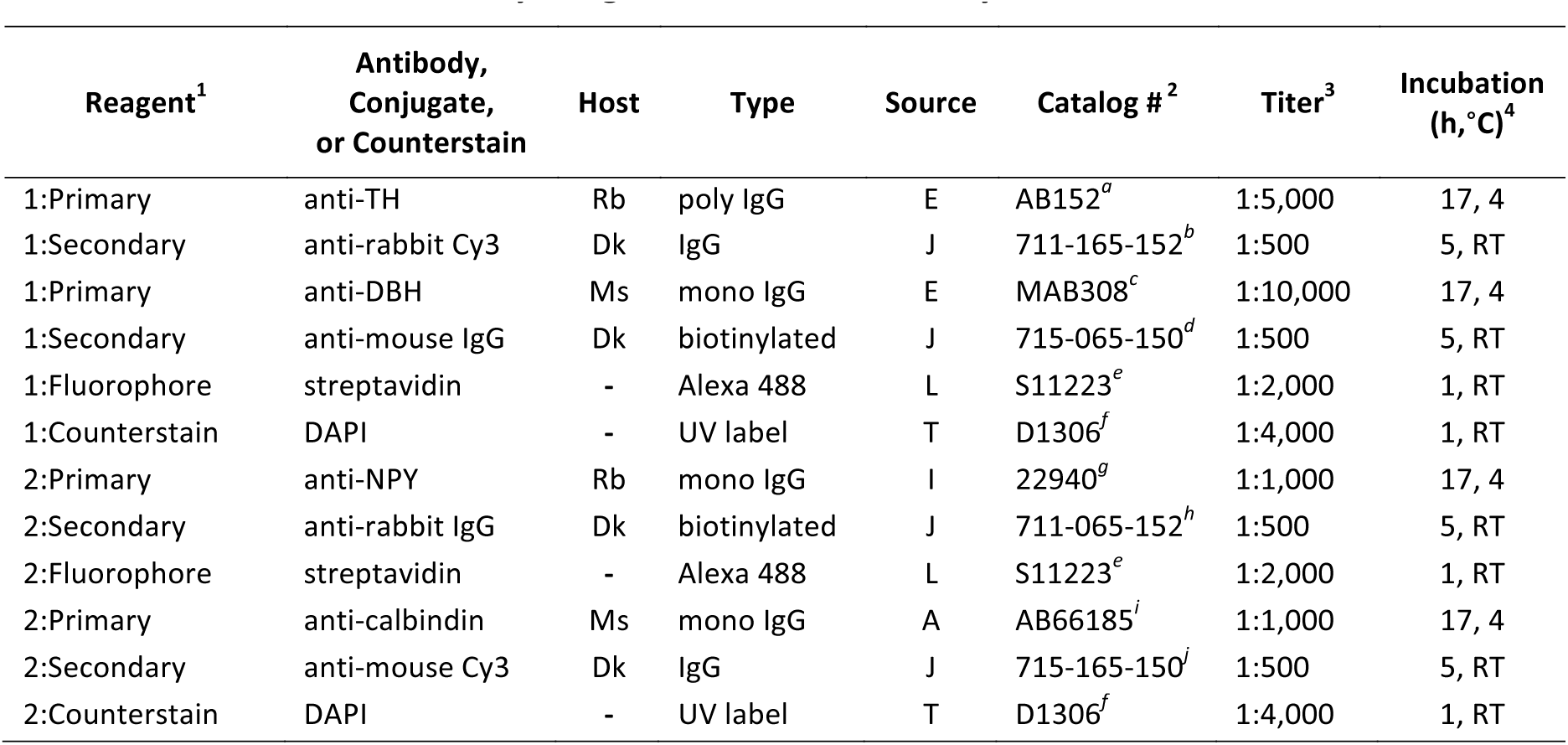
Immunohistochemistry Reagents Used in This Study

Tissues were processed for immunohistochemistry as described previously [28-31]. Briefly, all sections were placed for 1.5 hr in a blocking solution consisting of normal donkey serum (2%; EMD Millipore; Catalog #S30-100ML, Lot #2510142), Triton X-100 (0.1%; Sigma-Aldrich; Catalog #T8532-500ML, Lot #MKBH4307V) and TBS (pH 7.4 at room temperature). With the exception of selected tissue sections where the primary antibody step was omitted (*w/o 1°*), sections were then removed from blocking solution and incubated in a cocktail of primary antibodies according to the reaction parameters summarized in **Table 4**. All primary and secondary antibodies were prepared in blocking solution. After five washes in TBS, each for five min (5 × 5), all sections (including *w/o 1°*) were then incubated in a cocktail of secondary antibodies (**Table 4**). All sections were washed again (5 × 5) in TBS, reacted with fluorophore conjugates also prepared in blocking solution, and counterstained (**Table 4**). Following another 5 × 5 rinse in TBS, sections were mounted onto Superfrost slides and coverslipped with sodium bicarbonate-buffered glycerol (pH 8.6 at room temperature) and sealed with clear nail polish.

### 2.4.10 Photomicrography and post-acquisiton image processing of immunostained tissues

Immunostained and *w/o 1°* sections were visualized with the appropriate filters under epifluorescence illumination using the same microscope and software (and were photographed and images exported in the same manner) as described in *Section 2.4.6* for the Nissl-stained sections. Files of both individual and merged channels were imported into Adobe Photoshop, linearly adjusted for brightness and contrast across experiments, cropped, and saved as lossless TIFF-formatted images.

### 2.5 Methods pertaining to tissue fixed by immersion

#### 2.5.1 Immersion fixaton

In two instances, during collection efforts at Location 2 (**Table 1**), animals were deeply sedated and their morphological measurements recorded as described in *Section 2.3.1*, but were then subjected to immersion fixation rather than transcardial perfusion. Specifically, fully anesthetized animals were manually decapitated in the field between the second and third cervical vertebrae using large surgical scissors. The heads were placed immediately in buffered formalin (bottled liquid fixative containing a final concentration of 4% formaldehyde; i.e., “Source 1” in *Section 2.3.1*).

#### 2.5.2 Diffusible iodine-based contrast-enhanced computed tomography (DiceCT)

The specimens immersion-fixed in the field (*Section 2.5.1*) were transferred into aqueous solutions of Lugol’s iodine (I_2_KI) at the Oklahoma State University Center for Health Sciences [25, 26]. Specimen UTEP 21389 (*Rhampholeon boulengeri*) (trans-quadratic width (TQW) = 8.1 mm) was fully submerged in a 3.75% (i.e., 1.25% w/v of I_2_ and 2.5% w/v of KI) for seven days without replacement (of Lugol’s iodine solution). Specimen UTEP 21388 *(Trioceros johnstoni)* (TQW = 15.5 mm) was fully submerged in a 5.25% (i.e., 1.75% w/v of I_2_ and 3.5% w/v of KI) for 21 days without replacement. Upon immersion in Lugol’s iodine, both specimens were agitated for 60 sec using a vortex mixer to remove air bubbles and ensure that the contrast agent accessed the deepest openings within the specimen (e.g., the endocranial space). Stained specimens were removed from the Lugol’s solution, rinsed with tap water to remove excess stain, blotted dry, sealed individually in polyethylene bags to prevent dehydration, and then loaded into plastic mounting units for scanning.

Specimens were μCT-scanned at the Microscopy and Imaging Facility at the American Museum of Natural History (New York, NY), with a 2010 GE phoenix v|tome|x s240 high-resolution microfocus computed tomography system (General Electric, Fairfield, CT). A standard X-ray scout image was obtained prior to scanning to confirm specimen orientation and define the scan volume. The specimen UTEP 21389 was μCT-scanned at 130 kV and 150 μA for approximately 45 min and the specimen UTEP 21388 was μCT-scanned at 140 kV and 180 μA for approximately 90 min. Both specimens were imaged using a 0.1 mm copper filter, a tungsten target, air as the background medium, and 400 ms X-ray exposure timing with 3-times multi-frame averaging. Both were scanned at isometric voxel sizes (at 16.185 and 29.476 μm, respectively), and slices were assembled on an HP z800 workstation (Hewlett-Packard, Palo Alto, CA) running VG Studio Max (Volume Graphics GmbH, Heidelberg, Germany).

## 3. Results

### 3.1 Fixaton and storage

*Field-perfused specimens*. Five lizards were perfusion-fixed under similar conditions in the field (**Table 1**, **Fixation Type = ‘P’**). The field-perfusion procedure, from initial sedative exposure to brain storage, averaged 47.8 min (range 33-58 min), including 32 min (range 21-39 min) from exsanguination to brain storage. The brains remained unfrozen in storage solution (*Section 2.4.3a*) for an average of 46.2 days (range 21-54 days) (**Table 1**).

*Laboratory-perfused specimens*. Three lizards were perfused in the UTEP Biodiversity Collections Laboratory. These laboratory-based perfusions, from initial sedative exposure to brain storage (for these cases, on ice), averaged 37.7 min (range 29-50 min) (**Table 2**).

*Immersion-fixed specimens*. For the two lizards whose brains were immersion-fixed in the field (**Table 1**, **Fixation Type = ‘I’**), the process, from initial sedative exposure to immersion in buffered formalin (*Section 2.5.1*), averaged 21.5 min (range 18-25 min), which included time for full sedation and for morphometric measurements to be performed.

### 3.2 Qualitative evaluaton of tissue integrity and cytoarchitecture of perfusion-fixed tissues

With the unaided eye, surface vessels and sinuses free of blood, together with a uniformly pale, opaque color (**Figs. 1E-G**), indicated successful formaldehyde infusion and fixation of the brain under field conditions. This observation was corroborated at the light microscope level, where we observed that qualitatively the Nissl staining for the field-perfused animals was robust (**Fig. 2**) and appeared comparable, in terms of color richness and stain evenness, to stained sections obtained from brains perfused under our standard laboratory protocol (**Fig. 3**; see *Section 2.4.1b* and [28]). Specific brain regions and landmarks were delimited easily in the stained sections under both field and lab perfusion fixation conditions (**Figs. 2** and **3**). **Figure 4** shows a field-perfused agamid, the only specimen perfused with the fixative containing 10% formaldehyde, displaying tissue quality that was comparable to the field-caught specimens perfused with fixative solutions containing 4% formaldehyde. Similar to these latter specimens, the agamid displayed clearly defined cytoarchitectural features, including discernible laminated structures (e.g., cortex medialis, *C m*, **Fig. 4A**) and white matter tracts (e.g., optic tract, Op tr, **Fig. 4B**). Both largely acellular (**Fig. 4D**, *Layer 2*) and cell-dense layers (**Fig. 4D**, *Layers 3/4*), with observable Nissl substance in neuronal perikarya, were visible. The results show qualitatively that field fixation using solutions containing 4% formaldehyde produce tissue that is fixed sufficiently for obtaining excellent preservation of brain cytoarchitecture.

**Figure 2.**
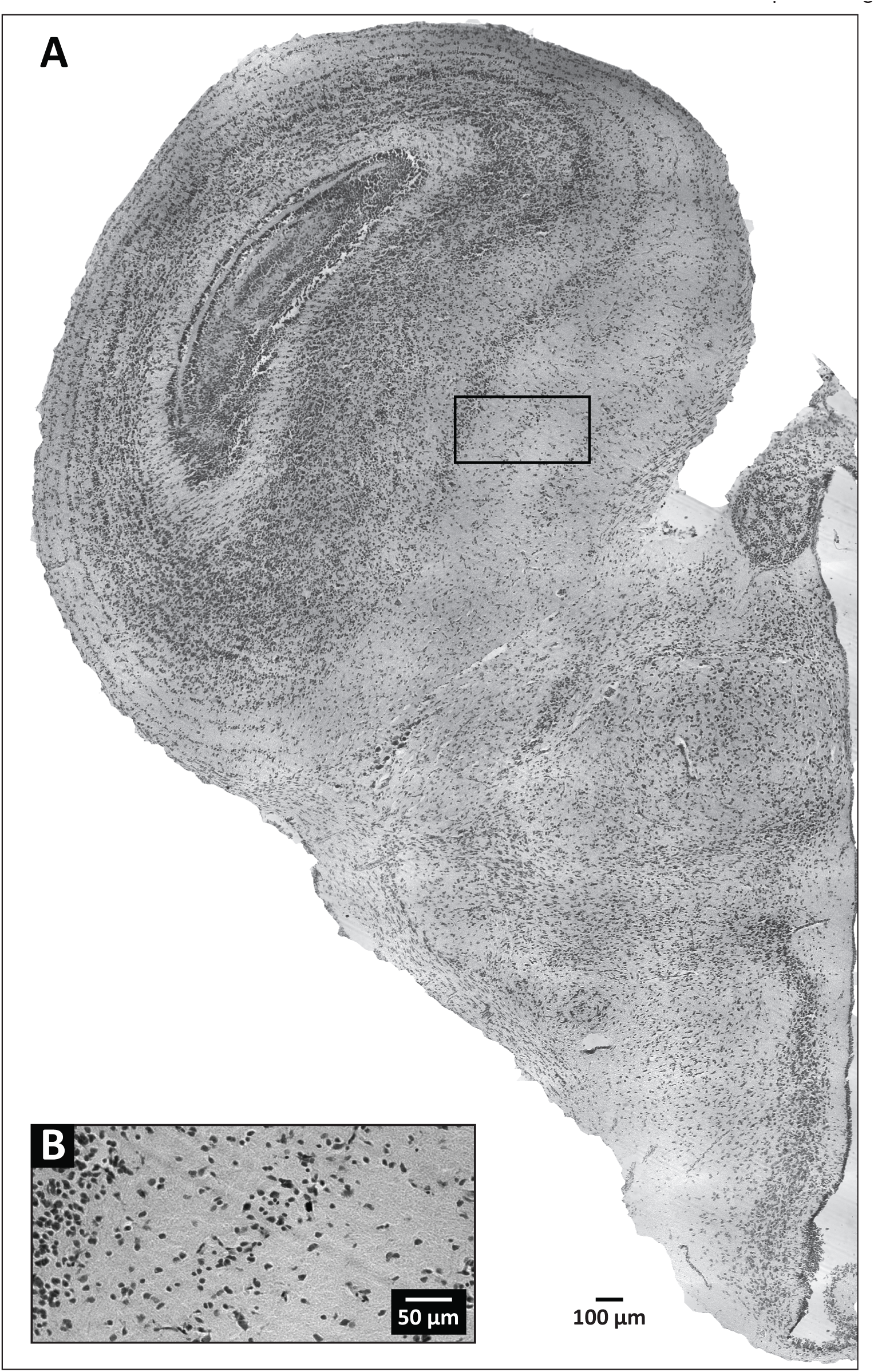
Representative tissue section at the level of the optic tectum, obtained from a field-perfused specimen of *Trioceros johnstoni*. (A) Wide field image of a hemisphere from a section of the specimen. The black box outlines the area enlarged in (B), which provides details regarding the level of background staining and cellular labeling demonstrable by our Nissl-based staining procedure.

**Figure 3.**
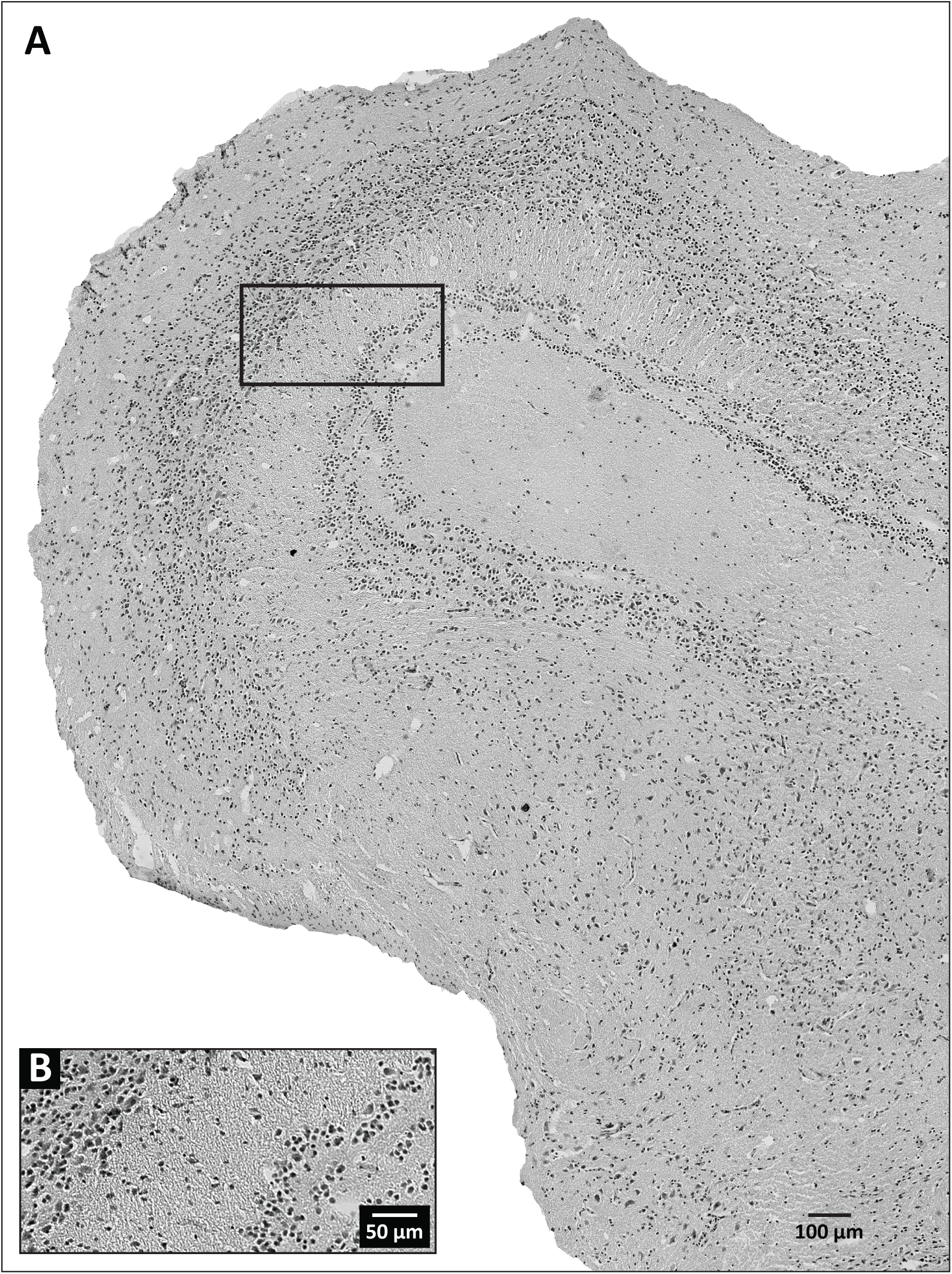
Representative tissue section at the level of the optic tectum, obtained from a laboratory-perfused specimen of *Trioceros jacksonii*. (A) Wide field image of a hemisphere from a section of the specimen. The black box outlines the area enlarged in (B), which provides details regarding the level of background staining and cellular labeling demonstrable by our Nissl-based staining procedure.

**Figure 4.**
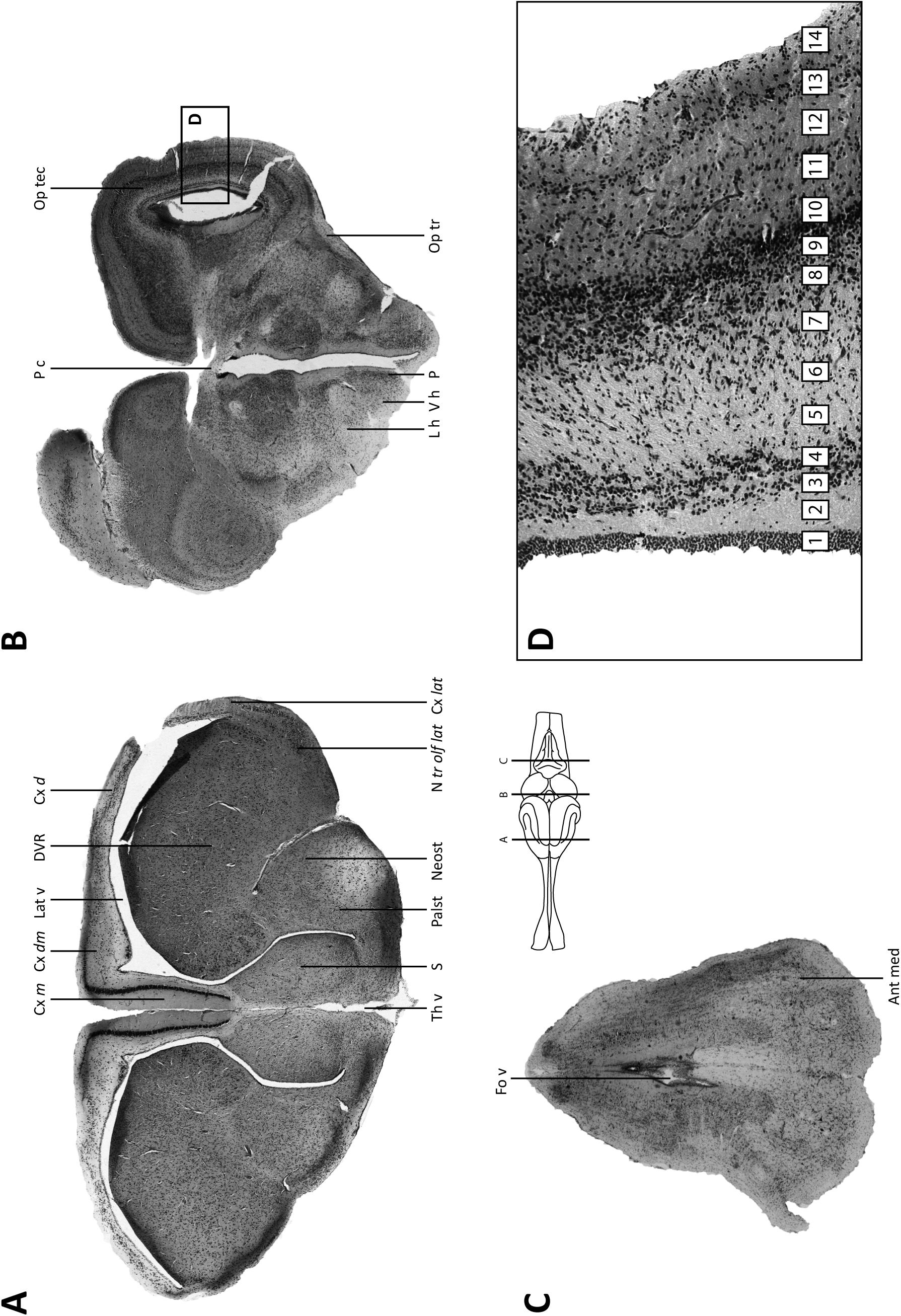
Photomicrographs of Nissl-stained brain sections from an agamid lizard. (A-C) Major brain regions are represented (A-forebrain; B-midbrain; C-hindbrain). (D) Detailed image of the optic tectum. The brain schematic was adapted from a drawing of a lizard rendered by artist Christiaan van Huijzen for Poster 2 of the poster book accompanying [40]. Delineation of major brain regions (A-C) generally follows [41-49] with only cosmetic changes made to the abbreviation style. The laminar organization of the optic tectum (D) follows [50]. Abbreviations: Ant med-Anterior medulla; Cx *d*-Cortex dorsalis (dorsal cortex); Cx *dm*-Cortex dorsomedialis (dorsomedial cortex); Cx *lat*-Cortex lateralis (lateral cortex); Cx *m*-Cortex medialis (medial cortex); DVR-Dorsal Ventricular Ridge; Fo v-Fourth ventricle; L h-Lateral hypothalamus; Lat v-Lateral ventricle; Neost-Neostriatum; N *tr olf lat*-Nucleus tractus olfactori lateralis (nucleus of the lateral olfactory tract); Op tr-Optic tract; Op tec-Optic tectum; Palst-Paleostriatum; P-Periventricular hypothalamus; P c-Posterior commissure; S-septal nuclei; Th v-Third ventricle; V h-Ventral hypothalamus.

### 3.3 Semi-quantitative evaluation of tissue integrity and cytoarchitecture of tissues perfusion-fixed using solutions containing 4% formaldehyde

The raw scored data obtained from the tissue evaluations of three independent raters (see **Supporting File S1**) are represented as a heat map in **Figure 5**. The map clearly shows distinct patterns between the two perfusion methods. For example, inter-observer variability was high, but intra-observer variability across criteria was generally not. In some cases, all observers agreed well on the conditions of the tissue (**Fig. 5**, *see black box outlines)*. **Table 5** shows the GEE analysis results for the correlated ordinal response based on the reduced mode scores (see **Supporting File S2** for the original R script). The analyses based on the raw data yielded similar conclusions (data not shown). Specifically, no significant difference was found between perfusion methods for criteria A and B (i.e., degree of blood in brain [*P* = 0.59] and evenness of stain [*P* = 0.63]). We did detect significant differences between perfusion methods for the remaining four criteria (C-F), with p-values varying from 0.0386 to < 0.0001 (**Table 5**). More specifically, tissue sections obtained from field perfusions received generally higher scores for these criteria, as suggested by the positive signs of the beta estimates and patterns readily deducible in the heat map (**Fig. 5**).

**Figure 5.**
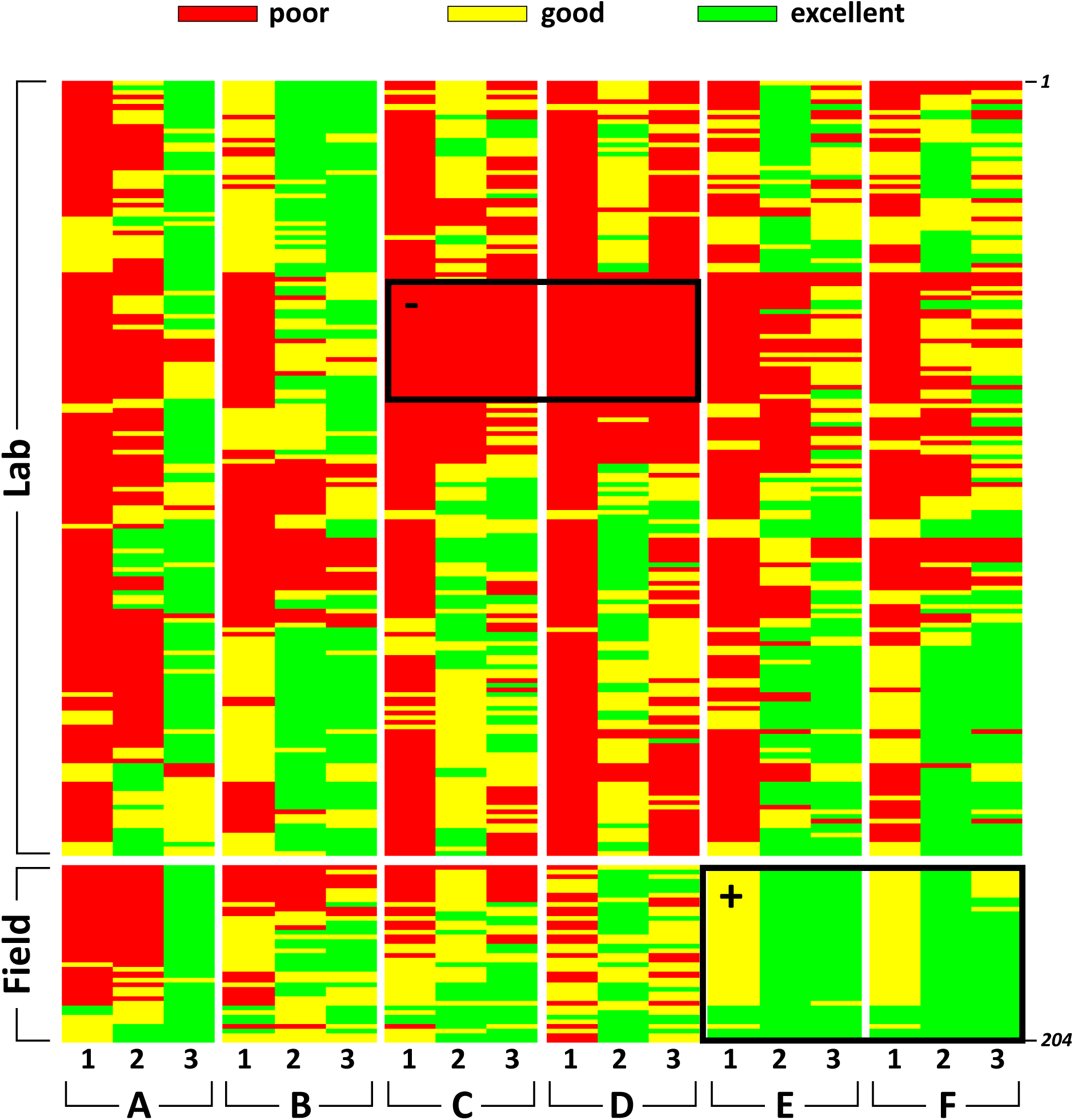
Heat map of scored semi-quantitative data for six qualitative variables from three independent observers. Observers evaluated Nissl-stained tissue sections (n = 204) from laboratory and field treatments using solutions containing 4% formaldehyde. Each column indicated with a small number (1, 2, or 3) represents an observer. Columns are grouped according to the qualitative variable being rated: presence of blood in tissue (A); evenness of stain (B); integrity of tissue at center of section (C); integrity of tissue at edges of section (D); clarity of lamination paterns (E); visibility of cell areas and nuclei (F). Tissue sections prepared under laboratory conditions are positioned on top (n = 166) and those prepared under field conditions on bottom (n = 38). The color code for the scored data is shown above the heat map. The black box outlines denoted with a ‘-‘ or a ‘+’ indicate selected regions of tissue ratngs (negatve or positve, respectvely) that were largely uniform across observers.

**Table 5.** Results of GEE analyses of clustered ordinal scores for scored data from stained tissue sections comparing field and lab fixation treatments using solutions containing 4% formaldehyde.

| Variable | Description | beta Estimate | Robust SE | Z | P-Value |
| --- | --- | --- | --- | --- | --- |
| A | presence of blood | 0.376 | 0.702 | 0.536 | 0.5922 |
| B | evenness of stain | −0.375 | 0.771 | −0.487 | 0.6261 |
| C | tissue integrity at center | 1.579 | 0.763 | 2.068 | 0.0384 |
| D | tissue integrity at edge | 7.091 | 0.631 | 11.231 | 0.0000 |
| E | lamination patterns visible | 26.920 | 0.492 | 54.726 | 0.0000 |
| F | cell clusters and nuclei visible | 23.577 | 6.194 | 3.806 | 0.0001 |

Results of GEE analyses of clustered ordinal scores for scored data from stained tissue sections comparing field and laboratory fixation treatments using solutions containing 4% formaldehyde.

### 3.4 Immunohistochemical staining of perfusion-fixed tissue

The results of our immunohistochemical staining are presented in **Figure 6A-D**. Robust TH-immunoreactive (-ir) neurons were observed in the periventricular hypothalamus of tissues fixed under field conditions for *Trioceros johnstoni* (**Fig. 6A**, *white arrows*). Fine TH-ir neurites (many of them likely axonal) were also observed within this region (**Fig. 6A**, *solid yellow horizontal lines)*. Fluorescent counterstaining additionally revealed that delicate structures, such as the ependymal layer lining the third ventricle, were largely intact under field fixation conditions (**Fig. 6A**, *large arrowheads)*. Similarly, laboratory-fixed tissues revealed robust TH-ir neurons in the periventricular hypothalamus of *Rieppeleon kerstenii* (**Fig. 6B**), yet the ependymal layer lining the third ventricle did not remain entirely intact and fine TH-ir neurites were less prominently visible. Additionally, NPY-labeled neurons and/or axonal fibers were localized readily in the field-fixed tissues of *Rhampholeon boulengeri* (**Fig. 6C**), and the laboratory-fixed tissues of *Rieppeleon kerstenii* (**Fig. 6D**). In contrast to these observations, tissue sections processed through all reaction steps but in the absence of primary antibody did not display any specific staining (see insets: **Fig. 6a-d**).

**Figure 6.**
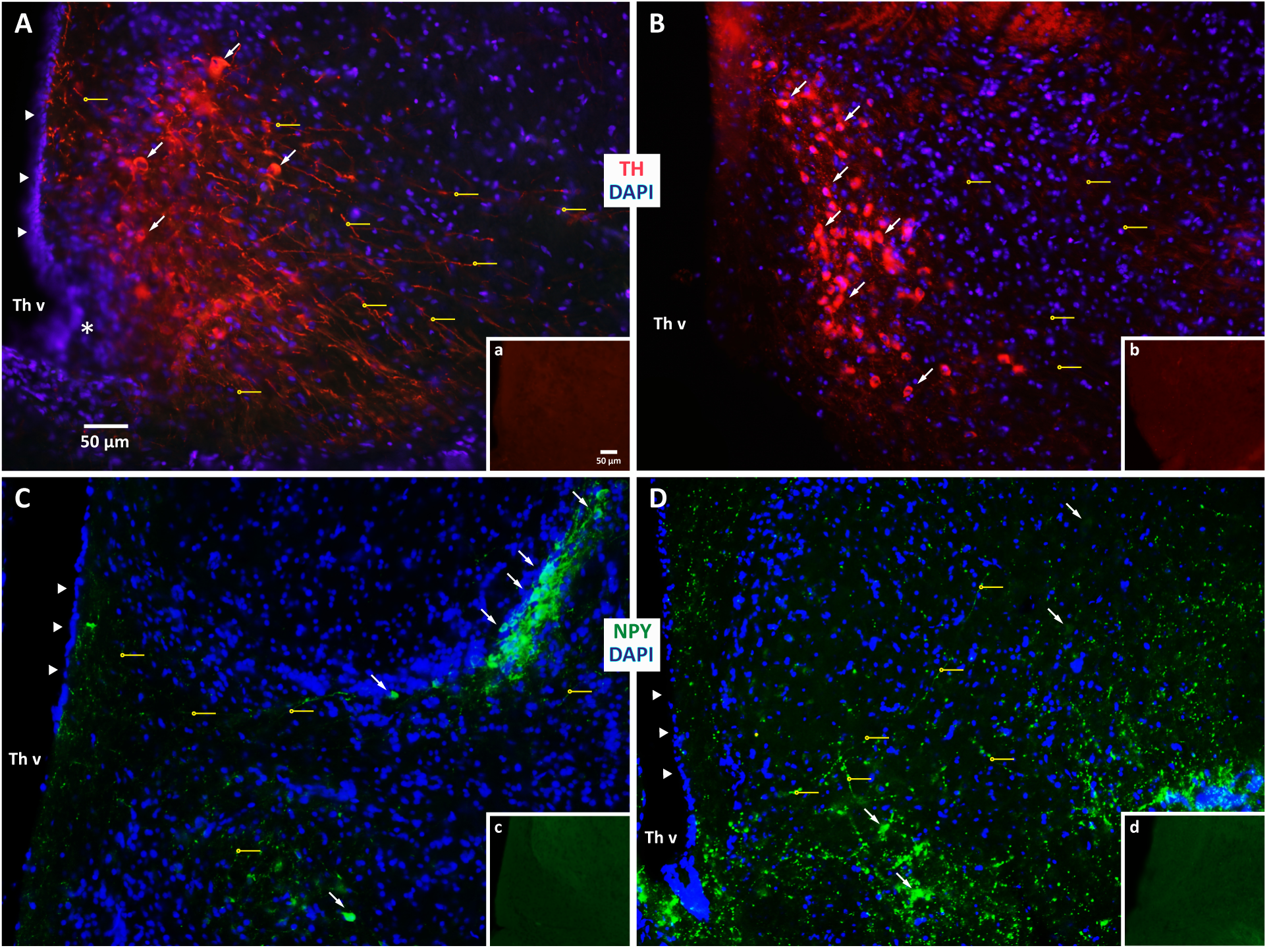
Comparison of immunohistochemical staining of brain tissue fixed under field and laboratory conditons. (A, B). The images show tyrosine hydroxylase immunoreactvity (-ir) (TH; *red*) with DAPI fluorescent counterstain (*blue*) for (A) *Trioceros johnstoni* fixed under field conditons, and (B) *Rieppeleon kerstenii* fixed under laboratory conditons. (C, D). The images show neuropeptde Y-ir (NPY; *green*), again with DAPI (*blue*) for (C) *Rhampholeon boulengeri* fixed under field conditons and (D) *Rieppeleon kerstenii* fixed under laboratory conditons (note that tissues in B and D are from the same animal). Both immunoreactve neurons (*arrows*) and neuronal extensions (*small solid horizontal lines ending in hollow circles*) are clearly visible, many of the later being identfiable axons with varicosites. The ependymal cell layers lining the third ventricle (Th v) in A, C and D are indicated by arrowheads. The single-plane image in A rendered a portion of the image slightly out of focus (*asterisk*). Insets (a–d) show views of sections processed in the absence of the primary antibody. For inset b, the image has been brightened linearly so that the tissue section can be clearly seen in the photo. Scale bars (panel A, inset a) apply to all remaining panels and insets, respectvely.

### 3.5 Immersion-fixed samples scanned using difusible iodine-based contrast-enhanced computed tomography (diceCT)

Two lizards (*Rhampholeon boulengeri* and *Trioceros johnstoni*) were prepared for diceCT scanning under the field conditions described in *Section 2.5.1*. In addition to bony tissues that are typically captured with X-ray imaging techniques, our contrast-enhanced specimens revealed, in great detail, differentiation between muscular, epithelial, glandular, and neural tissues (**Fig. 7**). This was equally true for superficial structures (e.g., hyobranchial muscles, distal branches of peripheral nerves) as it was for tissues located more deeply within the head, such as the brain. Indeed, even the internal anatomy of the brain was visualized clearly owing to the ability for Lugol’s iodine to differentiate between myelinated and non-myelinated components of the central (and peripheral) nervous systems (compare, for example, optic tectum [Op tec] and the optic tract [Op tr] in **Fig. 7C**). The high levels of contrast for these scans make them amenable to successful 3-D reconstruction of the soft anatomy (e.g., the brain and its peripheral cranial nerves), registration with the surrounding skull, and finer, Nissl-based visualization methods.

**Figure 7.**
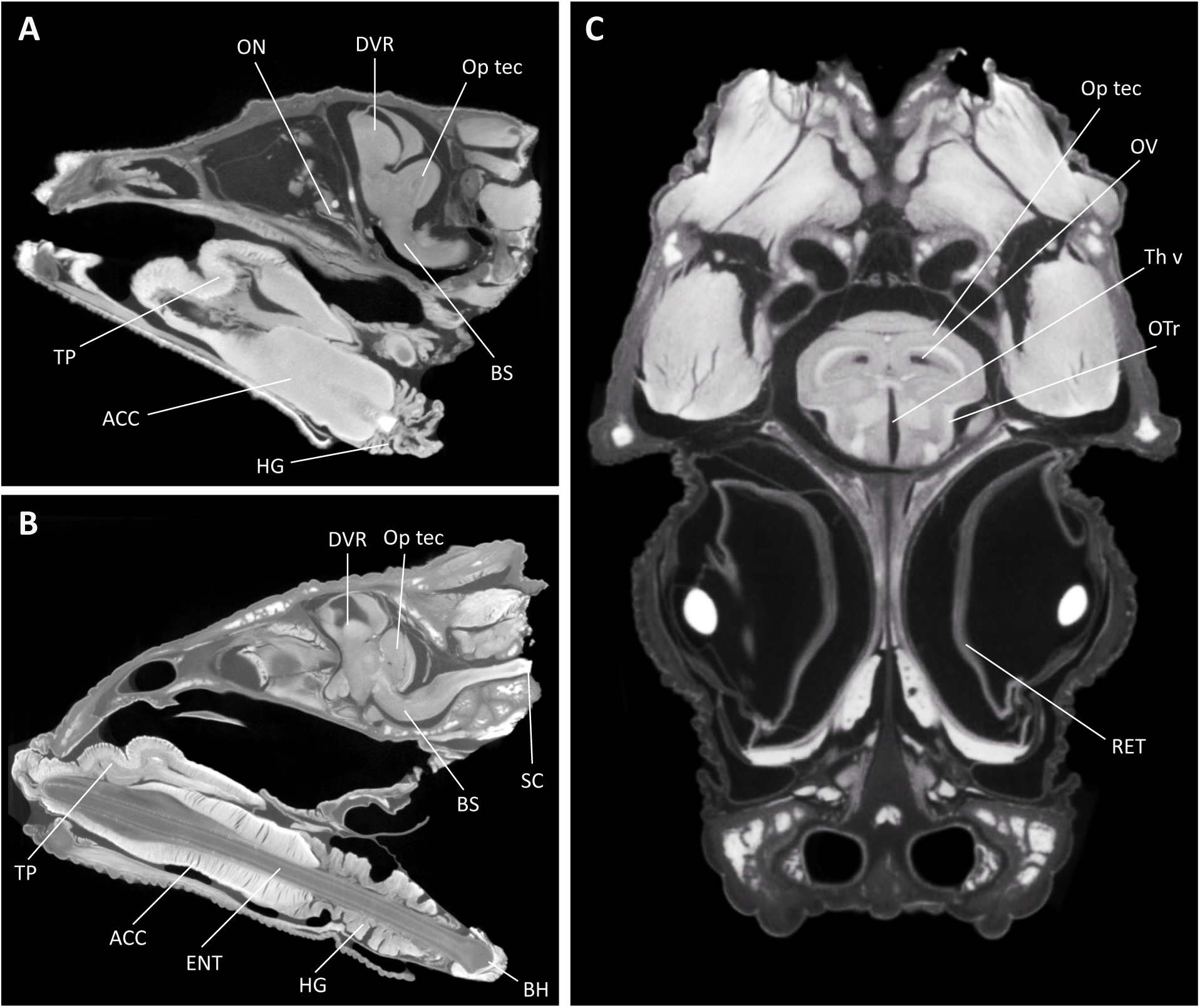
Difusible iodine-based contrast-enhanced computed tomography (DiceCT) through the heads of two chameleon species. (A) Parasagittal view of an adult male representative of *Rhampholeon boulengeri*; (B) parasagittal and (C) frontal views of an adult female representative of *Trioceros johnstoni*. These images illustrate the extraordinary diversity of soft anatomical structures that can be clearly visualized with our approach, including myelinated and unmyelinated components of the brain. Abbreviations for selected structures: ACC – M. accelerator linguae; BH – basihyoid; BS – brain stem; DVR – dorsal ventricular ridge; ENT – entoglossal process; HG – M. hyoglossus; ON – optic nerve; Op tec – Optic tectum; OTr – olfactory tract; OV – optic ventricle; RET – retina; SC – spinal cord; Th v – third ventricle; TP – tongue pad.

## 4. Discussion

In this study, we have evaluated two standard laboratory brain fixation methods for use in remote field locations where resources are limited and environmental conditions for tissue preservation are suboptimal. First, we found that transcardial perfusion-based methods to preserve lizard brain tissue, performed in parts of a remote biodiversity hotspot, are comparable to laboratory-based use of these methods in maintaining tissue quality at the cellular level. This conclusion is based on careful validation of the field fixation methods against laboratory methods by semi-quantitative cytoarchitectonic analysis of the processed brain tissue and by indirect immunohistofluorescence cytochemistry. Second, we found that immersion fixation in the field preserves gross neuroanatomical features of the brain very well, as evaluated using diceCT imaging. Moreover, the visualization of myelinated and unmyelinated components of the brain using diceCT imaging supports the efficacy of our field immersion fixation approach. To our knowledge, this paper is the first published account of a successful attempt to perform and rigorously validate protocols for transcardial perfusion and immersion fixation of brain tissue in a completely mobile field setting.

### 4.1 Methodological considerations

We found that field-perfused lizard brains are similar in condition to lizard brains perfused under standard laboratory conditions. In particular, cytoarchitectural features normally found in Nissl-stained brain sections were evident within the field specimens we processed, including discrete nuclear boundaries and structurally intact patterns of lamination. Moreover, the general appearance of the tissue, cleared completely of any blood, indicated complete perfusion. Whereas traditional methods to preserve brain tissue involve perfusing animals with ice-cold fixative solutions [51, 52], our collected animals were perfused with fixative solutions that were exposed to environmental conditions produced in the rainy season of a humid tropical climate, where we had no access to ice or cold storage. Specifically, our field-collected samples were subjected to high ambient temperatures (33-37°C) and a wide range of climates during their shipment from Central Africa to the southwestern United States. Considering the potential damage to delicate brain tissues from exposure to environmental variables and warm fixative solutions, our observations of no demonstrable differences between field-and lab-processed tissues for the first two criteria we evaluated (presence of blood and evenness of staining) is somewhat surprising. However, it has been shown that varying formaldehyde concentration within fixative solutions has little effect on the size of nuclei over a 10-fold range (1-20%), and extreme changes have only been observed in tissues fixed in at least 40% formaldehyde [53]. Furthermore, tissue shrinkage was not evident with the naked eye, which is known to occur in tissues incompletely fixed in formaldehyde or those subjected to varying temperatures [53].

The greatest negative effect on tissue quality for Nissl staining is arguably the interval between the time *post mortem* and the time of fixation [54]. It has been observed that fixation within 10 *h post mortem* has no effect on the intensity of stains, yet the ability for the tissue to be stained gradually deteriorates as the time interval increases, until staining capacity is entirely lost if fixation occurs 60-72 *h post mortem* [55]. On average, our field-collected animals were fixed in under an hour. Therefore, the mean duration between the time *post mortem* and the time of fixation for our field procedure does not compromise tissue quality. However, to minimize operational time, adequate training and multiple practice runs in a controlled environment are warranted before trying this procedure in the field.

#### 4.1.1 Cytoarchitectonics

We used a semi-quantitative approach to evaluate the cytoarchitecture of the tissue sets processed under laboratory and field conditions of perfusion fixation. A recent survey of semiquantitative methods used to evaluate histology has found that there is no accepted standard for the types of criteria used for such evaluations [56]. Given the many diverse approaches used to rate the quality of brain tissue, the study recommended that at the very least, investigators should provide a rationale for the specific criteria they use [56]. In line with this recommendation, we note here that the criteria we used were selected on the basis of the goals of our larger experimental research program involving these species, which are to examine the gross neuroanatomical relationships among their gray and white matter structures and the general cytoarchitectonic features of their brain tissue such as aggregations of neurons forming nuclei and laminae. These scales of comparative analysis are informed, in part, by seminal comparative neuroanatomical studies published at the turn of the twentieth century by Ramón [50], Edinger [57], and Brodmann [58]. Our approach necessarily constrains the criteria we use to those listed in *Section 2.4.7*, guided as we are by a rationale of general histological evaluation at the tissue level rather than its examination at the single-cell or subcellular levels. If the focus were on more fine-grained studies of the morphology or intracellular structure of the cells (e.g., the appearance of neurites, the condition of the organelles), then the criteria chosen using Nissl-based methods would likely be different (e.g., see Chapters X-XIV of Barker [59]), and alternatives to the Nissl method would also have been considered (e.g., see Chapter II in Vol. I of Cajal [60]).

Our statistical analyses show that, on the basis of the two major criteria we used to evaluate fixation efficacy (presence of blood and evenness of stain), no differences in tissue quality were observed between field-and laboratory-perfused specimens. This result supports our qualitative observations of the tissue samples. Further, on the basis of the remaining four criteria (integrity of tissue in center, integrity of tissue at edges, visualization of lamination patterns, visualization of cell clustering and nuclei), the field-perfused tissue apparently displayed significantly better tissue quality than lab-perfused tissue. While these results may be somewhat surprising given the more controllable fixation conditions generally available in a laboratory setting, we must interpret these findings with caution for a few reasons. First, from a qualitative standpoint, it is difficult to separate these four criteria from underlying effects that could be due to other confounding factors, such as tissue damage that was incurred during the mounting of the tissue sections onto glass slides, tissue adherence to the slides during their mechanical transfer through separate reagent reservoirs during the thionin staining procedure, and any differences in tissue stability resulting from variations in section thickness or in the pH and salt composition of the fixative solutions we used. Second, our analysis is limited by the variability we observed among the three independent raters, despite the fact that the mixed model we used to analyze our results mitigates this issue to some extent. Finally, these latter four criteria are interdependent to a large degree on the first two criteria; this interdependence of predictors makes it possible that the statistically significant effects we obtained may be more apparent than real, given that the first two major criteria (which are not dependent, or as dependent, on issues such as mounting and mechanical transfer) show no differences between the two groups.

#### 4.1.2 Immunohistochemistry

In addition to providing validation of our fixation procedures in the field by evaluating cytoarchitectonic criteria within Nissl-stained tissue sections, we sought indications that our fixation was compatible with standard chemoarchitectural localization methods. In particular, given that the field conditions required prolonged post-fixation in formalin-sucrose, we were concerned about the possibilities of over-fixing the tissue. The duration of formaldehyde fixation may lead to absent or weak binding for some epitopes, preventing effective chemoarchitectural studies with immunohistochemistry [61]. At times, poor penetration of the fixative can occur with too short of an exposure time and excessive cross-linkage can occur from prolonged exposure [62]. In addition, prolonged formalin fixation can cause irreversible damage to some epitopes, but this is largely dependent on the antibody used [63].

For our immunohistochemical procedures, we first aimed to identify dopamine-containing neurons in the periventricular hypothalamus, a well-studied neuronal subpopulation that has been documented previously to be present within the lizard brain [64–66]. Our results demonstrating robust TH-immunoreactivity (-ir) in neurons of field-fixed tissues that is comparable to that observed in laboratory-fixed tissues, confirms the findings of others [64–66] that have characterized this cell population as dopaminergic and extends them by demonstrating that the field conditions of prolonged post-fixation did not prevent chemical identification of these neurons. Relative to the field-fixed sample, the lab-fixed sample displayed apparently elevated levels of TH expression in cell bodies and low levels in fibers. Whether this staining difference indicates a difference in species, fixation efficacy, or peptide transport from the cell bodies to distal neurites between animals, is unclear. We also found comparable labeling of NPY-ir, reported to be present in chameleon brain [67], in cell bodies and/or axonal fibers within field-and lab-fixed tissues. These findings demonstrate the extensibility of our field-based fixation methods to antigens of different types (i.e., those that mark the presence of small neurotransmitters or those that mark neuropeptides). Although immunohistochemical labeling and visualization was feasible using our approach, further research is required to understand the degree of immunohistochemical reactivity across a broader range of antigens within field-perfused brains under our protocol conditions.

#### 4.1.3 DiceCT imaging

In addition to cytoarchitectonic and immunohistochemical validation of our procedures in the field, diceCT scans from immersion-fixed field-collected samples show that our field protocol is compatible with the non-destructive visualization of gross neuroanatomical features in relation to other soft-tissue structures of the head as well as the skull. Our approach to prepare specimens for diceCT scans, immersion fixation and prolonged storage in fixative followed by iodine-enhancement, was demonstrated to have no negative effects on the contrast and visual quality of the scans. Gray and white matter regions were clearly distinguishable in the soft tissue, further demonstrating that our specimen preparation was successful. Importantly, diceCT can be achieved without encephalectomy, allowing for the interrelationships between central and peripheral components of the nervous system to be preserved. Another advantage of diceCT is that 3-D rendering software can be used to rapidly visualize the high-resolution scans as complex, 3-D soft-tissue anatomy [25, 26]. In turn, these datasets can be analyzed to quantify and compare neuroanatomical structures among different body regions and species [68]. Indeed, the possibility with field-fixed samples to visualize neural circuitry from the cellular level to that of entire brain regions sets up the potential for a comprehensive mapping of brain interconnectedness in three dimensions across multiple scales.

#### 4.1.4 Summary remarks about methodology

In sum, both our qualitative and semi-quantitative comparisons of field-and laboratory-perfused brain samples indicate that none of several variables that differed between laboratory and field experiments (post-fixation duration, formalin pH and buffer composition, environmental exposure) was sufficient to produce major differences in tissue quality, staining or visualization. Moreover, immersion fixation of samples in the field produced tissue that was readily compatible with non-destructive diceCT imaging. Further studies are needed to determine whether laboratory-based immersion fixation with less environmental exposure produces comparable tissue quality for these species that is amenable to diceCT imaging. We anticipate this to be the case given the evidence provided by other lab-based studies [25, 26].

### 4.2 Considerations in the field

Access and/or availability of supplies are the major limiting factors for field research, especially for long-term expeditions in remote locations with poor infrastructure. Biodiversity hotspots [2] are distributed disproportionally in tropical countries, which have generally high levels of poverty [69]. Most of the equipment and supplies detailed herein should be procured before travel, as they may not be available in certain countries, especially underdeveloped ones. Our expedition was no different from other carefully planned herpetological collecting expeditions in which it was realized in retrospect that certain supplies should have been brought to the field [70, 71]. For example, including pH test strips (pH paper) or a battery-powered pH meter in our field kit would have allowed for us to make accurate pH measures of our fixatives, and we recommend researchers to include this item before embarking on their own collecting trips. Indeed, our list of supplies was far from complete, and can also be modified to facilitate more specialized perfusion approaches intending to address specific research questions. With respect to essential supplies, particular attention should be paid to formaldehyde, a hazardous material that is not permitted on commercial airlines. Researchers attempting this procedure in underdeveloped countries or countries without reliable access to formaldehyde (at a range of concentrations) must ensure that this material can be acquired upon their arrival as it will be a major limiting factor. Typical sources of formaldehyde include universities, morgues, or laboratory supply companies. Researchers should also bear in mind the disposal of chemical waste generated from this procedure, which takes the form of a very small amount of waste formalin (˜2 ml/perfusion). We recommend researchers attempting this procedure to store any amount of hazardous waste in a labeled container and dispose of this waste at a proper facility when one is made available.

To accomplish this procedure under completely mobile conditions in the field over a relatively long period of time without access to a laboratory, both a substantial amount of fluids and replacements of most supplies are required. Ideally, a researcher would have access to some sort of basecamp with basic shelter from the elements, where most of the supplies can be stored. In this case, animals can be captured and transported to basecamp for the fixation procedure, which will likely cut down on waste, sample exposure to environmental conditions, and potential equipment loss.

Perfusion fixation of brain tissue has been performed in various forms for at least the better part of a century [e.g., 72, 73]. Clearly, our methodology is only one approach to conduct perfusion fixation of brain tissue. Pump-assisted perfusions, for example, can also be used. Electric pumps are optimal because both the hydrostatic pressure and flow rate of the fixative solutions are controllable [74, 75]. However, pumps require a reliable source of electricity, which is often not available in the field. Gravity assisted approaches are also a viable alternative to transcardial perfusions [76] and are arguably superior to syringes for controlling the pressure of injected solutions. However, it can be challenging to bring an entire gravity-fed perfusion system (e.g., containers, fluid lines) when working in very remote locations; yet, a resourceful scientist can utilize a variety of common items to replace containers, such as water bottles. Nevertheless, if the containers are too small, it can be difficult to properly regulate the flow rate of solutions in the fluid lines as they are drained from their elevated positions (E. D. Roth, personal communication). Immersion fixation is arguably the easiest fixation method to achieve in the field. This approach can help avoid inflicting physical damage to the brain tissue during dissection. However, it is widely recognized that transcardial perfusion fixation produces superior results to immersion fixation with respect to staining efficacy and visible immunoreactivity [77–80]. Finally, our procedure works well for small lizards; however, larger animals require greater amounts of fixative to penetrate deep brain tissues and a small syringe will likely not suffice. This problem should be recognized before an expedition is undertaken and can be easily resolved with the use of a larger syringe or gravity-assisted perfusion set-up with a sizable elevated container. For very large animals where such a set-up is not feasible, some unique brain fixation methods in the field have been described [81, 82].

### 4.3 Concluding remarks

We have shown that field-based brain fixation methods can preserve effectively the cytoarchitecture, chemoarchitecture, and gross neuroanatomy of the brains of wild-caught herpetofauna. Specifically, performing transcardial perfusion fixation in a mobile field setting is an advantageous alternative to laboratory-based perfusion. The tissue integrity and stain intensity obtained in this study indicate, qualitatively and semi-quantitatively, that the degree of environmental exposure and amount of formaldehyde concentration did not negatively impact our visualization of neural substrates within the tissues. Also, we found that immersion fixation of the intact brain and skull is highly feasible for preserving gross anatomy and for preparing specimens for diceCT imaging. Collectively, these protocols should serve as a flexible framework for researchers attempting field-based fixation of brain tissue. Our approach also has the potential to liberate researchers from laboratory limitations imposed by traditional methods and can be harnessed to explore species diversity that is critically needed for neuroscientific and gross anatomical research.

Accelerated declines in global biodiversity are associated inescapably with losses in unknown amounts of trait variation. We must endeavor to mitigate these losses with higher rates of rescue for as many types of data as possible. Our understanding of the anatomical diversity of the vertebrate brain is rudimentary; for this reason, there recently have been efforts to increase the level of species diversity in neuroscientific research [83]. Although novel ways to salvage existing brain specimens will undoubtedly help in this effort [5], this goal will best be accomplished by the continued active collection of brain specimens from wild animals in the field.

## 5. Acknowledgments

For their companionship and assistance in the field, we are grateful to: Dr. Mathias Behangana, Lukwago Wilber, Dr. Chifundera Kusamba, Mwenebatu M. Aristote, Wandege M. Muninga, and Jean-Pierre Mokanse Watse. Dr. Baluku Bajope of the Centre de Recherche en Sciences Naturelles (CRSN) provided project support and permits, and the Institut Congolais pour la Conservation de la Nature (ICCN) kindly granted permits to work in protected areas in DRC. We thank the Uganda Wildlife Authority (UWA) of Kampala for necessary permits to work in Uganda, and Mr. James Lutalo, Commissioner of Wildlife Conservation and CITES Authority for Uganda. We are also grateful to Dr. Xiaogang Su, Director of the BBRC Statistical Consulting Laboratory (SCL), for assistance with the statistical analyses used in this study. The SCL is supported by grant 5G12MD007592 from the National Institute on Minority Health and Health Disparities (NIMHD), a component of the National Institutes of Health (NIH).

For support throughout this project, we owe special thanks to all members of the UTEP Systems Neuroscience Laboratory, especially Berenise De Haro. We also thank Morgan Hill and Henry Towbin of the American Museum of Natural History’s Microscopy and Imaging Facility for assistance. EG and DFH are supported by grant DEB-1145459 from the National Science Foundation (NSF) of the United States. Work in the UTEP Systems Neuroscience Laboratory is supported by NIH grant GM109817 awarded to AMK and a Grand Challenges Grant awarded to AMK by the UTEP Office of Research and Sponsored Projects. Additionally, research work by AMK and EG is supported by a Howard Hughes Medical Institute STEM Education Grant (UTEP PERSIST; awarded to AMK and EG as co-PIs; PI = Dr. Stephen Aley). DFH and EMW are supported by Keelung Hong Graduate Fellowships and AM is supported by an NSF STEM Fellows in K-12 Education (GK-12) Fellowship. PMG is supported by grants DEB-1457180 and EAGER 1450850 from NSF and the Oklahoma State University Center for Health Sciences Department of Anatomy and Cell Biology.

## 6. Author Contributions

*Experiment conception and design:* AMK, DFH. *Specimen acquisition:* DFH, EG. *Field perfusions:* DFH. *Laboratory perfusions:* DFH, CSL, EMW. *Nissl staining:* DFH, EMW. *Immunohistochemistry:* EMW, DFH. *Microscopy, wide-field imaging:* EMW, *DFH. Microcomputed tomography:* PMG. *Data analysis:* DFH, AMK, EMW, AM, KN. *Reagents/materials/analysis tools:* AMK, EG, PMG, CSL. *Manuscript preparation:* AMK, DFH with contributions from EG, PMG, EMW.

## 8. Supporting Information

*File S1*. Raw scores used for statistical analysis, MS Excel-formatted file.

*File S2:* Original R script used for the statistical analysis, MS Word-formatted file.

## References

1. Mora C, Tittensor DP, Adl S, Simpson AGB, Worm B. How many species are there on Earth and in the ocean? PLoSBiology 2011; 9: e1001127.

2. Mittermeier RA, Turner WA, Larsen FW, Brooks TM, Gascon C. Global biodiversity conservation: The critical role of hotspots. In: Zachos FE, Habel JC, editors. Biodiversity Hotspots: Distribution and Protection of Conservation Priority Areas. Berlin: Springer-Verlag; 2011. p. 3–22.

3. Myers N. Biodiversity hotspots revisited. Bioscience 2003; 53: 916–917.

4. Hoffman M, Hilton-Taylor C, Angulo A, Böhm M, Brooks TM, Butchart SHM et al. The impact of conservation on the status of the world’s vertebrates. Science. 2010; 230: 1503–1509.

5. Iwaniuk AN. The importance of scientific collecting and natural history museums for comparative neuroanatomy. Annals of the New York Academy of Sciences. 2011; 1225 (Suppl 1): E1–E19.

6. Rocha LA, Aleixo A, Allen G, Almeda F, Baldwin CC, Barclay MV et al. Specimen collection: An essential tool. Science. 2014; 344: 814–815.

7. Lieb CS, Buth DG, Gorman GC. Genetic differentiation in *Anolis sagrai*: A comparison of Cuban and introduced Florida populations. Journal of Herpetology. 1983; 17: 90–94.

8. Kolbe JJ, Glor RE, Schettino LR, Lara AC, Larson A, Losos JB. Genetic variation increases during biological invasion by a Cuban lizard. Nature. 2004; 431: 177–181.

9. Stuart YE, Campbell TS, Hohenlohe PA, Reynolds RG, Revell LJ, Losos JB. Rapid evolution of a native species following invasion by a congener. Science. 2014; 346: 463–466.

10. Kamath A, Stuart YE, Campbell TS. Behavioral partitioning by the native lizard *Anolis carolinensis* in the presence and absence of the invasive Anolis sagrei in Florida. Breviora. 2013; 535: 1–10.

11. Kamath A, Stuart YE. Movement rates of the lizard *Anolis carolinensis* (Squamata: Dactyloidae) in the presence and absence of *Anolis sagrei* (Squamata: Dactyloidae). Breviora. 2015; 546: 1–7.

12. Huelsenbeck JP, Bull JJ, Cunningham CW. Combining data in phylogenetic analysis. Trends in Ecology and Evolution. 1996; 11: 152–158.

13. Levasseur C, Lapointe F-J. War and peace in phylogenetics: A rejoinder on total evidence and consensus. Systems Biology. 2001; 50: 881–891.

14. Gonzalez-Voyer A, Winberg S, Kolm N. Brain structure evolution in a basal vertebrate clade: Evidence from phylogenetic comparative analysis of cichlid fishes. BMC Evolutionary Biology. 2009; 9: 238.

15. Charvet CJ, Sandoval AL, Striedter GF. Phylogenetic origins of early alterations in brain region proportions. Brain, Behavior & Evolution. 2010; 75: 104–110.

16. Sylvester JB, Rich CA, Loh YE, van Staaden MJ, Fraser GJ, Streelman JT. Brain diversity evolves via differences in patterning. Proceedings of the National Academy of Sciences of the United States of America. 2010; 107: 9718–9723.

17. Carlson BA, Hasan SM, Hollmann M, Miller DB, Harmon LJ, Arnegard ME. Brain evolution triggers increased diversification of electric fishes. Science. 2011. 332: 583–586.

18. Balanoff AM, Bever GS, Rowe TB, Norell MA. Evolutionary origins of the avian brain. Nature. 2013; 501: 93–96.

19. Albert FW, Sommel M, Carneiro M, Aximu-Petri A, Halbwax M, Thalmann O et al. A comparison of brain gene expression levels in domesticated and wild animals. PLoS Genetics. 2012; 8: e1002962.

20. Editorial. Worldwide neuroscience. Nature Neuroscience. 2003; 6: 901.

21. Yusuf S, Baden T, Prieto-Godino LL. Bridging the gap: Establishing the necessary infrastructure and knowledge for teaching and research in neuroscience in Africa. Metabolic Brain Disease. 2014; 29: 217–220.

22. Hughes DF, Gignac PM, Lieb CS, Greenbaum E, Khan AM. Fixing to collect? A validated brain tissue preservation method for mitigating the loss of neuroanatomical data in threatened biodiversity hotspots. [Program #429.04] 2015 Abstract Viewer/Itinerary Planner. Washington, DC: Society for Neuroscience; 2015. Online.

23. Mittermeier RA, Gil PR, Hoffman M, Pilgrim J, Brooks T, Mittermeier CG et al. Hotspots Revisited: Earth’s Biologically Richest and Most Endangered Terrestrial Ecoregions. Mexico City, Mexico: CEMEX; 2004.

24. Plumptre AJ, Davenport TRB, Behangana M, Kityo R, Eilu G, Ssegawa P et al. The biodiversity of the Albertine Rift. Biological Conservation. 2007; 134: 178–194.

25. Gignac PM, Kley NJ. Iodine-enhanced micro-CT imaging: Methodological refinements for the study of the soft-tissue anatomy of post-embryonic vertebrates. Journal of Experimental Zoology. Part B, Molecular and Developmental Evolution. 2014; 322: 166–176.

26. Gignac PM, Kley NJ, Clarke JA, Colbert MW, Morhardt AC, Cerio D et al. Diffusible iodine-based contrast-enhanced computed tomography (diceCT): An emerging tool for rapid, high-resolution, 3-D imaging of metazoan soft tissues. Journal of Anatomy. In press.

27. Berod A, Hartman BK, Pujol JF. Importance of fixation in immunohistochemistry: Use of formaldehyde solutions at variable pH for the localization of tyrosine hydroxylase. Journal of Histochemistry and Cytochemistry. 1981; 29: 844–850.

28. Khan AM, Watts AG. Intravenous 2-deoxy-D-glucose injection rapidly elevates levels of the phosphorylated forms of p44/42 mitogen-activated protein kinases (extracellularly regulated kinases 1/2) in rat hypothalamic parvicellular paraventricular neurons. Endocrinology. 2004; 145:351–359.

29. Khan AM, Ponzio TA, Sanchez-Watts G, Stanley BG, Hatton GI, Watts AG. Catecholaminergic control of mitogen-activated protein kinase signaling in paraventricular neuroendocrine neurons in vivo and in vitro: A proposed role during glycemic challenges. Journal of Neuroscience. 2007; 27: 7344–7360.

30. Khan AM, Kaminski KL, Sanchez-Watts G, Ponzio TA, Kuzmiski JB, Bains JS et al. MAP kinases couple hindbrain-derived catecholamine signals to hypothalamic adrenocortical control mechanisms during glycemia-related challenges. Journal of Neuroscience. 2011; 31: 18479–18491.

31. Khan AM, Walker EM, Dominguez N, Watts AG. Neural input is critical for arcuate hypothalamic neurons to mount intracellular signaling responses to systemic insulin and deoxyglucose challenges in male rats: Implications for communication within feeding and metabolic control networks. Endocrinology. 2014; 155: 405–416.

32. Watson RE, Wiegand SJ, Clough RW, Hoffman GE. Use of cryoprotectant to maintain long-term peptide immunoreactivity and tissue morphology. Peptides. 1986; 7: 155–159.

33. Lenhossék M. Der Feinere BauDesNervensystems imLichte Neuester Forsuchungen. 2nd ed. Berlin: H. Kornfield; 1895.

34. Kiernan JA. Classification and naming of dyes, stains and fluorochromes. Biotechnic & Histochemistry. 2001; 76: 261–278.

35. R Core Team. R: A language and environment for statistical computing. Vienna, Austria: R Foundation for Statistical Computing; 2015. Available: https://www.R-project.org/.

36. Heagerty PJ, Zeger SL. Marginal regression models for clustered ordinal measurements. Journal of the American Statistical Association. 1996; 91: 1024–1036.

37. Halekoh U, Højsgaard S, Yan J. The R Package geepack for Generalized Estimating Equations. Journal of Statistical Software. 2006; 15(2). Available: http://www.jstatsoft.org/.

38. Morona R, González A. Calbindin-D28k and calretinin expression in the forebrain of anuran and urodele amphibians: Further support for newly identified subdivisions. Journal of Comparative Neurology. 2008; 511: 187–220.

39. Moreno N, Domínguez L, Morona R, González A. Subdivisions of the turtle Pseudemys scripta hypothalamus based on the expression of regulatory genes and neuronal markers. Journal of Comparative Neurology. 2012; 520: 453–478.

40. Nieuwenhuys R, Ten Donkelaar HJ, Nicholson C. The Central Nervous System of Vertebrates. Berlin: Springer; 1998.

41. Foster RE, Hall WC. The connections and laminar organization of the optic tectum in a reptile (*Iguana iguana*). Journal of Comparative Neurology. 1975; 163: 397–425.

42. Northcutt RG. Forebrain and midbrain organization in lizards and its phylogenetic significance. In: Greenberg N, MacLean PD, editors. Behavior and Neurology of Lizards. Bethesda: National Institutes of Mental Health; 1978. p. 11–64.

43. Shanklin WM. The central nervous system of *Chameleon vulgaris*. Acta Zoologica. 1930; 11:425–490.

44. Senn DG, Northcutt RG. The forebrain and midbrain of some squamates and their bearing on the origin of snakes. Journal of Morphology. 1973; 140: 135–152.

45. Smeets WJAJ, Hoogland PV, Lohman AHM. A forebrain atlas of the lizard *Gekko gecko*. Journal of Comparative Neurology. 1986; 254: 1–19.

46. Greenberg N. A forebrain atlas and stereotaxic technique for the lizard, *Anolis carolinensis*. Journal of Morphology. 1982; 174: 217–236.

47. Northcutt RG. Architectonic studies of the telencephalon of *Iguana iguana*. Journal of Comparative Neurology. 1967; 130: 109–147.

48. Butler AB, Northcutt RG. Retinal projections in *Iguana iguana* and *Anolis carolinensis*. Brain Research. 1971; 26: 1–13.

49. Butler AB, Northcutt RG. Architectonic studies of the diencephalon of *Iguana iguana* (Linnaeus). Journal of Comparative Neurology. 1973; 149: 439–461.

50. Ramón P. El structura de encéfalo del cameléon [Structure of the chameleon brain]. Revista TrimestralMicrográfca. 1896; 1: 146–182. Spanish.

51. Aitken PG, Breese GR, Dudek FF, Edwards F, Espanol MT, Larkman PM et al. Preparative methods for brain slices: A discussion. Journal of Neuroscience Methods. 1995; 59: 139–149.

52. Lipton P, Aitken PG, Dudek FE, Eskessen K, Espanol MT, Ferchmin PA et al. Making the best of brain slices: Comparing preparative methods. Journal of Neuroscience Methods. 1995; 59: 151–156.

53. Fox CH, Johnson FB, Whiting J, Roller PP. Formaldehyde fixation. Journal of Histochemistry and Cytochemistry. 1985; 33: 845–853.

54. Scudamore CL, Hodgson HK, Patterson L, Macdonald A, Brown F, Smith KC. The effect of post-mortem delay on immunohistochemical labelling—a short review. Comparative Clinical Pathology. 2011; 20: 95–101.

55. Gu J, Huang WM, Polak JM. Stability of immunocytochemical reactivity of neuronal substances following delayed fixation. Journal of Neuroscience Methods. 1985; 12: 297–302.

56. Klopfeisch R. Multiparametric and semiquantitative scoring systems for the evaluation of mouse model histopathology-a systematic review. BMC Veterinary Research. 2013; 9: 123.

57. Edinger L. Untersuchungen über die vergleichende Anatomie des Gehirnes. 4. Studien über das Zwischenhirn der Reptilien [Studies on the comparative anatomy of the brain. 4. Studies on the hypothalamus of reptiles]. Abhandlungen der Senckenbergischen Naturforschenden Gesellschaft. [Treatises of the Senckenberg Nature Research Society]. 1899; 20: 21–197 (with three color plates following). German.

58. Brodmann K. *Vergleichende Lokalisationslehre der Groβhirnrinde in ihren Prinzipien dargestellt auf Grund des Zellenbaues*. Leipzig: Johann Ambrosius Barth; 1909. German. Translated by Laurence J. Garey as Brodmann’s Localisation in the Cerebral Cortex. The Principles of Comparative Localisation in the Cerebral Cortex Based on Cytoarchitectonics. *3rd ed*. New York: Springer Science + Business Media; 2006.

59. Barker LF. The Nervous System and its Constituent Neurones. New York: D. Appleton and Co; 1899.

60. Cajal SR. *Histology of the Nervous System of Man and Vertebrates*. Translated by Neely Swanson and Larry W. Swanson. New York: Oxford University Press; 2 vols.; 1995. Originally published as Histologie du Système Nerveux de L’homme et des Vertébrés (2 volumes). Translated by L. Azoulay Paris: Maloine; 2 vols.; 1909, 1911.

61. Srinivasan M, Sedmak D, Jewell S. Effect of fixatives and tissue processing on the content and integrity of nucleic acids. American Journal of Pathology. 2002; 161: 1961–1971.

62. Werner M, Chott A, Fabiano A, Battifora H. Effect of formalin tissue fixation and processing on immunohistochemistry. American Journal of Surgical Pathology. 2000; 24: 1016–1019.

63. Wasielewski RV, Werner M, Nolte M, Wilkens L, Georgii A. Effects of antigen retrieval by microwave heating in formalin-fixed tissue sections on a broad panel of antibodies. Histochemistry. 1994; 102: 165–172.

64. Bennis M, Calas A, Geffard M, Gamrani H. Distribution of dopamine immunoreactive systems in brain stem and spinal cord of the chameleon. Biological Structures and Morphogenesis. 1990-1991; 3: 13–19.

65. González A, Smeets WJAJ. Catecholamine systems in the CNS of amphibians. In: Smeets WJAJ, Reiner A, editors. Phylogeny and Development of Catecholamine Systems in the CNS of Vertebrates. Cambridge: Cambridge University Press; 1994. p. 77–102.

66. Smeets WJAJ. Catecholamine systems in the CNS of reptiles: Structure and functional correlations. In: Smeets WJAJ, Reiner A, editors. Phylogeny and Development ofCatecholamine Systems in the CNS of Vertebrates. Cambridge: Cambridge University Press; 1994. p. 103–134.

67. Bennis M, Bam’hamed S, Rio JP, Le Cren D, Repérant J, Ward R. The distribution of NPY-like immunoreactivity in the chameleon brain. Anatomy and Embryology. 2001; 203: 121–128.

68. Gold MEL, Shulz D, Budassi M, Vaska P, Gignac PM, Norell MA. Flying starlings and PET lend insights on the evolution of volant dinosaurs. Current Biology. In press.

69. Jackson K. Mean and Lowly Things: Snakes, Science, and Survival in the Congo. Cambridge, MA: Harvard University Press; 2008.

70. James J. The Snake Charmer: A Life and Death in Pursuit of Knowledge. New York: Hyperion; 2008.

71. Sachs JD, Mellinger AD, Gallup JL. The geography of poverty and wealth. Scientific American. 2001; 284: 70–75.

72. Carmichael EA. Microglia: An experimental study in rabbits after intracerebral injection of blood. Journal of Neurology and Psychopathology. 1929; 9: 209–216.

73. Koenig H, Groat RA, Windle WF. A physiological approach to perfusion-fixation of tissues with formalin. Biotechnic & Histochemistry. 1945; 20: 13–22.

74. Eichhammer P, Zeller R, Rohkamm R. Fixation of neural tissue for electron microscopy with an electronically controlled perfusion pump. Tissue and Cell. 1987; 19: 153–157.

75. Hoops D. A perfusion protocol for lizards, including a method for brain removal. MethodsX. 2015; 2: 165–173.

76. Rieke GK, Bowers DE, Silvy NJ. A technique for improved fixation of the pigeon central nervous system for electron microscopy. Anatomical Record. 1981; 200: 121–125.

77. Gertz SD, Rennels ML, Forbes MS, Nelson E. Preparation of vascular endothelium for scanning electron microscopy: A comparison of the effects of perfusion and immersion fixation. Journal of Microscopy. 1975; 105: 309–313.

78. Tago H, Kimura H, Maeda T. Visualization of detailed acetylcholinesterase fiber and neuron staining in rat brain by a sensitive histochemical procedure. Journal of Histochemistry and Cytochemistry. 1986; 34: 1431–1438.

79. Beach TG, Tago H, Nagai T, Kimura H, McGeer PL, McGeer EG. Perfusion-fixation of the human brain for immunohistochemistry: Comparison with immersion-fixation. Journal of Neuroscience Methods. 1987; 19: 83–192.

80. Bondonna TJ, Jacquet Y, Wolf G. Perfusion-fixation procedure for immediate histologic processing of brain tissue. Physiology & Behavior. 1977; 19: 345–347.

81. Knudsen SK, Mørk S, Øen EO. A novel method for in situ fixation of whale brains. Journal of Neuroscience Methods. 2002; 120: 35–44.

82. Manger PR, Pillay P, Maseko BC, Bhagwandin A, Gravett N, Moon D-J et al. Acquisition of brains from the African elephant (*Loxodonta africana*): Perfusion-fixation and dissection. Journal of Neuroscience Methods. 2009; 179: 16–21.

83. Striedter GF, Belgard TG, Chen C-C, Davis FP, Finlay BL, Güntürkün O et al. NSF workshop report: Discovering general principles of nervous system organization by comparing brain maps across species. Journal of Comparative Neurology. 2014; 522: 1445–1453.

